# Fast phylogenetic generalised linear mixed-effects modelling using the glmmTMB R package

**DOI:** 10.64898/2025.12.20.695312

**Authors:** Coralie Williams, Maeve McGillycuddy, Szymon M. Drobniak, Benjamin M. Bolker, David I. Warton, Shinichi Nakagawa

## Abstract

Phylogenetic generalised linear mixed models (PGLMMs) help ecologists to distinguish ecological drivers from other processes shaping evolutionary patterns, yet existing implementations are often limited in distributional scope or computational speed. We compare five R packages for fitting PGLMMs and highlight the new covariance structure `propto` in the general-purpose GLMM package `glmmTMB`. Simulations show that `glmmTMB` fits PGLMMs faster overall than `brms`, `MCMCglmm`, `INLA`, and `phyr`, while producing similar model estimates. We present the first practical application of `glmmTMB` for fitting phylogenetic random effects using likelihood-based models that accommodate repeated measures, demonstrated through case studies of evolutionary trait data. By improving both speed and flexibility, `glmmTMB` broadens access to PGLMM and supports deeper insights into trait evolution and diversification.

## 1 Introduction

Diversity of life occurs within complex ecological communities influenced by evolutionary history. In fact, it is now widely recognised that patterns in species traits are partly determined by evolutionary ancestry (Garamszegi, 2014; Martins & Hansen, 1997). This understanding has driven the development of phylogenetic comparative methods (PCMs) as essential tools for exploring how evolutionary relationships influence species trait diversity over time. Originally proposed by Felsenstein, 1985, PCMs have since evolved into a comprehensive framework that accounts for the non-independence among species due to common evolutionary origins (Cornwallis & Griffin, 2024; Hernández et al., 2013). These methods allow researchers to partition the variance attributable to evolutionary history from that driven by ecological interactions, thereby providing deeper insights into species evolution and ecology.

Among the earliest PCMs, phylogenetically independent contrasts (PICs) were developed to assess trait values as statistically independent contrasts along a phylogeny, enabling valid inferential statistical tests between species traits (Felsenstein, 1985). Building on PICs, phylogenetic generalised least squares (PGLS) incorporated phylogenetic relatedness directly into regression models, allowing for multiple predictors and formal hypothesis testing under different evolutionary models (Grafen, 1989; Martins & Hansen, 1997). More recently, phylogenetic linear mixed models (PLMMs) and generalised linear mixed models (PGLMMs) have added further flexibility, accommodating discrete traits, repeated measures, multiple responses, and multiple sources of non-independence through additional random effect terms (Gallinat & Pearse, 2021; Hadfield & Nakagawa, 2010; Halliwell et al., 2025; Ives & Helmus, 2011; Mizuno et al., 2025). Together, these models have broadened the scope of comparative research, enabling researchers to examine ecological and evolutionary drivers of diversity.

General purpose implementations of PGLMMs started developing 15 years ago with `MCMCglmm` (Hadfield, 2010), the `binaryGLMM` function in the `ape` package (Ives & Helmus, 2011; Paradis & Schliep, 2019) and more recently through packages such as `brms` (Bürkner, 2017) and `phyr` (Ives et al., 2020) for example. As trait datasets grow rapidly in size and resolution, especially with the rise of global databases and citizen science contributions, aggregating data to species-level averages obscures important within-species variation (Mouquet et al., 2012; Nakagawa et al., 2017). PGLMMs can incorporate this variation by partitioning ecological and evolutionary components of trait variance, which supports more nuanced inferences about trait evolution and diversification (Goolsby, 2015; Opedal et al., 2023). However, current PGLMM implementations still face limitations. First, the frequentist (maximum likelihood) options that exist are limited to a small number of non-Gaussian distributions and covariance structures. Second, Bayesian frameworks can be slow and computationally intensive, particularly for complex models or when applied to large datasets. Finally, accessible and generalised tutorials with applications remain scarce. These limitations may have restricted the broader uptake of PGLMMs, despite their value in modelling complex trait data. Filling this gap, in both flexibility and computational efficiency, will help unlock the full potential of contemporary trait data to inform both evolutionary theory and conservation strategies (Arnold & Nunn, 2010; Gallinat & Pearse, 2021).

In this study, we evaluate the performance of PGLMMs implementations in R statistical software for evolutionary ecological research and highlight the novel `propto` (proportional) covariance implementation in the `glmmTMB` package (Kristensen & McGillycuddy, 2025; McGillycuddy, 2023), which can be used to fit PGLMMs. This widely used package was designed for general purpose fitting of mixed models, so it can handle a wide variety of different data types and study designs that might not otherwise be available in software designed specifically for phylogenetic analysis. Using simulations, we compare `glmmTMB` with four commonly used R packages for fitting PGLMMs. These include two Bayesian packages (`brms` and `MCMCglmm`), the integrated nested Laplace approximation (INLA), and the frequentist `phyr` package. We focus on PGLMMs with repeated observations per species and assessed models run time. To demonstrate the use of PGLMMs in practice using `glmmTMB`, we use real example trait datasets involving within-species measures. Our goal is to provide ecologists with practical guidance for fitting PGLMMs using `glmmTMB` and other available implementations that make full use of rich trait datasets, allowing for more accurate inference of evolutionary patterns and processes. This work supports the wider adoption of flexible phylogenetic models that can reveal biological signals that may be missed when species-level averages are used. To demonstrate these models in practice and provide guidance for users, we provide an online tutorial on fitting PGLMMs.

## 2 Methods

We registered our study protocol in January 2025 (Williams et al., 2025), detailing the methodological plan following the ADEMP-PreReg template provided in Siepe et al. (2024). We followed our registered plans, requiring only minor changes detailed in ‘Additions and deviations’. We reported our simulation items in line with recommended reporting guidance (Morris et al., 2019; Williams et al., 2024).

### 2.1 Generalized linear mixed models

We introduce phylogenetic generalized linear mixed models (PGLMMs) for data with repeated measurements per species by building up step–by–step from a basic generalised linear mixed model (GLMM). The response is assumed to follow a scalar-valued probability distribution with location (mean) parameter and, where relevant, an additional scale or dispersion parameter. The following models are fitted using either maximum–likelihood or Bayesian estimation frameworks (see Box 1).

We begin with a GLMM, where the response vector ***y*** (*n* × 1) has mean ***μ*** linked to the predictors by a function *g*(*·*). The linear predictor combines fixed and random effects:

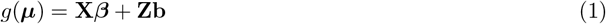

where **X** is the *n* × *p* matrix of fixed–effect predictors with coefficient vector ***β*, Z** is the *n* × *q* random–effect design matrix, and **b** ∼ 𝒩 (**0, Σ**) is the *q* × 1 vector of random effects. Conditional on **b**, the response follows a distribution ***y*** | **b** ∼ 𝒟 (***μ***, *ϕ*), where *ϕ* denotes a dispersion or residual variance parameter. In a standard GLMM, **Z, *b***, and **Σ** represent one or more independent random effects with unstructured covariance.

To incorporate evolutionary relatedness among species, we include a random effect whose covariance reflects phylogenetic relationships. Let **Z**_*p*_ map each observation to its species, and let 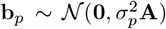 be the species–level random effects with variance 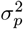 and where **A** is the phylogenetic correlation matrix derived from a phylogenetic tree. The linear predictor then becomes

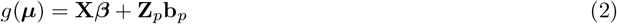

The phylogenetic correlation matrix is derived from an assumed model of trait evolution. The most common choice is the Brownian motion model, which assumes that trait values change gradually over time according to a random walk process along the branches of the phylogeny (Felsenstein, 1985; Martins & Hansen, 1997). Under this assumption, the expected variance in trait values between two species is proportional to the amount of shared evolutionary history, meaning that closely related species are more similar in their traits than distantly related ones. Alternative evolutionary models can be used to estimate macroevolutionary parameters (Blomberg et al., 2020), although these are more difficult to apply.

In many ecological datasets, multiple observations are recorded per species, for example repeated measurements across individuals, sites, or time points. To account for this structure, we extend the PGLMM to include two species–level random effects. The first captures phylogenetic covariance among species, while the second captures additional variation among species that is assumed independent of phylogeny. The linear predictor is then

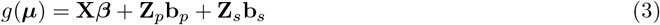

where **Z**_*p*_ and **Z**_*s*_ both map observations to species, 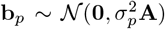 are the phylogenetically structured species effects, and 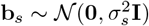 are the independent species effects with variance 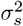.

The formulation in Equation 3 provides a flexible framework for partitioning total trait variance into phylogenetic, non-phylogenetic, and residual components (de Villemereuil et al., 2016; Hadfield & Nakagawa, 2010). This variance decomposition quantifies the relative contributions of evolutionary history and other ecological or biological factors to trait variation. Additional random effects can be included to account for other sources of variation, for example, temporal or spatial effects. Measures such as phylogenetic heritability (Housworth et al., 2004; Lynch, 1991), Pagel’s *λ* (Pagel, 1999) and Blomberg’s *K* (Blomberg et al., 2003) can be derived from these models and provide insight into the amount of phylogenetic signal and the processes shaping trait evolution (Pearse et al., 2025). For instance, 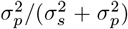 represents the portion of phylogenetic signal across all species effects in the model specification in Equation 3. When multiple observations per species are available, within-species variation can be explicitly estimated. If there is only one observation per species, the non-phylogenetic component (effect) is absorbed into the residual variance (i.e., potentially confounded by measurement errors). Notably, the non-phylogenetic effect, which may capture ecological influences on species similarities that are not explained by shared ancestry, stands in contrast to the phylogenetic effect that reflects shared evolutionary history.

`glmmTMB` can fit PGLMMs using the `propto` covariance structure, which specifies a multivariate random-effect that is proportional to a user-supplied phylogenetic correlation matrix. Further details on how to fit a PGLMM with `glmmTMB` are provided in our online supporting information.

### 2.2 Simulation study

#### 2.2.1 Model specification

We conducted a simulation study to evaluate run time and estimation accuracy of PGLMMs across five R packages (`phyr`, `glmmTMB`, `brms`, `MCMCglmm`, and `INLA`; see Box 1). We fitted models specified in Equation 3 across varying numbers of species *N*_*species*_ *∈* (25, 50, 100, 200, 400, 800), four different number of replicates per species (including balanced and unbalanced design; see below for details), and variance components, with two values each for both the independent species effect 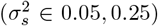 and the phylogenetic species effect 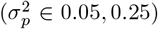. For datasets with *>*100 species we did not assess `brms` and `phyr` packages, as they were computationally intensive, requiring several days per model. We simulated data *N*_*sim*_ = 4,000 times for *≤*100 species and *N*_*sim*_ = 500 times for species *>*100. Across the combinations of parameters evaluated for Equation 3 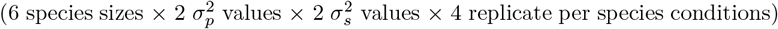, this resulted in a total of 216,000 simulations. Additionally, we evaluated models following the specification in Equation 2, representing datasets with a single measurement per species (*n*_reps_ = 1); results for these models are provided in Supporting Information. We assessed these models across the same number of species, same two values for the variance component 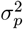, and numbers of simulation repetitions. We did not fit models with `phyr` for datasets containing only one measure per species as Equation 2 cannot be specified with this package. Full details of simulated parameters and the number of conditions assessed per model specification are provided in Supporting Information Table S2.

For the Bayesian models, we used the default specifications but increased the number of iterations, based on pilot runs, to ensure adequate convergence (see Supporting Information for details). Priors were set to convey only minimal information, representing the broad scale of the parameters while limiting their influence on estimation accuracy for the purpose of the simulation study. For `brms`, we used the package’s default priors, which are generally weakly informative and designed to provide regularisation while remaining flexible across a range of models. For `MCMCglmm`, we used the recommended parameter-expanded priors (see Supporting Information for more details).

#### 2.2.2 Data generation

The intercept coefficient was fixed to *β*_0_ = 1 and the slope coefficient *β*_1_ was set to 1.5. The continuous covariate *x* was generated from a uniform distribution ranging between 10 and 20 at the level of individual observation (non-species level) and was fixed for all simulation repetitions. The number of replicates per species was generated following both a balanced and an unbalanced design. For the balanced design, the number of replicates per species was fixed to *n*_*reps*_ *∈* (1, 5, 10, 30). For the unbalanced design, the number of replicates per species was generated from a beta distribution with *α* = 1.25 and *β* = 3, then multiplied by 99, rounded to the closest integer, and increased by 1. As a result, the number of observations per species ranged from 1 to 100, with an approximate mean of 30, median of 27, and mode of 12. For each simulation, a phylogenetic tree representing species evolutionary relationships was generated using the `rtree()` function from the ape R package (Paradis & Schliep, 2019), which simulates phylogenies through a recursive random splitting algorithm. Branch lengths of the simulated trees were assigned using the `compute.brlen()` function (Grafen, 1989), with the power parameter set to *α* = 1 to adjust the relative height of the tips. This produced ultrametric trees, from which we derived the phylogenetic correlation matrix **A** under a Brownian motion model of evolution. Specifically, **A** was computed using the `vcv()` function in the ape package, which yields a correlation structure corresponding to a linear decline of relatedness from the tips towards the root. The inverse correlation matrix of the phylogenetic relationships, for input in `MCMCglmm` and `INLA` models, was calculated using the `inverseA()` function from the `MCMCglmm` R package by inverting the phylogenetic correlation matrix (**A**). We ran simulations on R version 4.3.1 on the high-performance computing cluster ‘Katana’ supported by Research Technology Services at UNSW Sydney (UNSW, 2024).

#### 2.2.3 Performance metrics

We assessed the run time of models as the total time to compute, meaning for Bayesian models, this includes compilation and sampling time. We explored differences in compilation and sampling time for Bayesian packages using one dataset as a case study. We evaluated the simulated model estimates of the mean covariate with root mean squared error (RMSE), coverage and confidence interval width. We assessed the two random variance components (phylogenetic and non-phylogenetic) bias and RMSE. For the runtime and model estimate results, we excluded all models from a simulation run if any model failed to converge, ensuring fair comparison. We describe below and in the Supporting Information the convergence metrics used for each package in our simulation study.

#### 2.2.4 Additions and deviations

We deviated from the registered plan in how we handled non-converged cases. For the Bayesian packages (`MCMCglmm` and `brms`), rather than re-running models which did not achieve effective sample sizes of 400, we used package specific diagnostics and increased the number of iterations until at least 80% of the models reached a criterion. For `MCMCglmm`, the criterion was to pass the Heidelberger–Welch stationarity test for the fixed and random effect chains. For `brms` criteria were based on the bulk and tail effective sample sizes (ESS *≥* 400) and 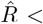 1.01 (Vehtari et al., 2021). We did not inspect diagnostic plots, as this was not feasible at the scale of the simulation study (but is recommended in practice). After preliminary testing we fixed package specific iteration counts for the entire study, setting `iter` = 20,000 for `brms` and `nitt` = 303,000 for `MCMCglmm`. In addition, we deviated from the protocol by scaling the covariate *x* to have mean zero and unit variance, to ensure that convergence issues could not be attributed to the predictor scale. For the `INLA` packages, non-convergence was flagged using the package built-in check (see Supporting Information for more detail). In addition, we labelled a model as non-converged if any random effect variance estimates exceeded 4, which in our settings is more than 16 times the true value. Convergence checks for the frequentist frameworks (`phyr` and `glmmTMB`) are detailed in the Supporting Information.

## 3 Results

### 3.1 Simulation study

Our simulation results show that `glmmTMB` converged in all but one of 216,000 simulations. Other frequentist packages, `phyr` converged in all (100%) simulation runs and `INLA` achieved convergence rates around 99% across all species sizes (Table 1). Convergence for `MCMCglmm` declined as the number of species increased, from 96.9% with 25 species to 67.3% at 800 species. `brms` converged in 79.0% to 89.8% of runs for 25 to 100 species (Table 1). The number of retained simulated models per package and the proportion retained after removing simulation runs that failed to converge are given in Table S3.

**Table 1:** Convergence rates percentage (%) of phylogenetic mixed models per package across *N*_species_. Results are based on 64,000 simulations for each *N*_species_ *≤* 100 and 8,000 simulations for each *N*_species_ *≥* 100, given varying variance components values (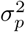 and 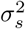), and varying number of replicates per species (*n*_rep_). Details of varying parameter values is provided in the Supporting Information Table S2.

| **Package** | $N_{\text{species}}$ | | | | | |
| --- | --- | --- | --- | --- | --- | --- |
|  | **25** | **50** | **100** | **200** | **400** | **800** |
| **brms** | 79.0 | 84.0 | 89.8 | – | – | – |
| **MCMCglmm** | 95.9 | 95.0 | 92.5 | 86.3 | 77.8 | 67.3 |
| **INLA** | 99.7 | 99.5 | 99.5 | 99.8 | 99.9 | 100 |
| **glmmTMB** | 100* | 100 | 100 | 100 | 100 | 100 |
| **phyr** | 100 | 100 | 100 | – | – | – |

\* one model did not converge.

From the simulations, we found that `glmmTMB`, was faster on average than the other packages across all dataset sizes, with relatively faster times as the number of species increased, except for 25 species models where `phyr` was slightly faster (Figure 1). The mean runtime of models across simulations increased rapidly with species number, most prominently for `phyr` and `MCMCglmm`, followed by other packages (Figure S1 and Table S4). For example, for 100 species, the mean runtime was 0.55 seconds for `glmmTMB`, compared with 3 minutes 42 seconds for `MCMCglmm` and 8 minutes 55 seconds for `brms`. The discrepancy became much larger at higher species numbers: for datasets with 400 species or more, `MCMCglmm` required over an hour (1 h 05 min for 400 species; 7 h 59 min for 800 species), whereas `glmmTMB` still finished within seconds (4.90 s and 25.92 s, respectively). For the Bayesian models, we assessed one simulated dataset and provide the breakdown of compilation and sampling time in the Supporting Information section 2.1.2.

**Figure 1:**
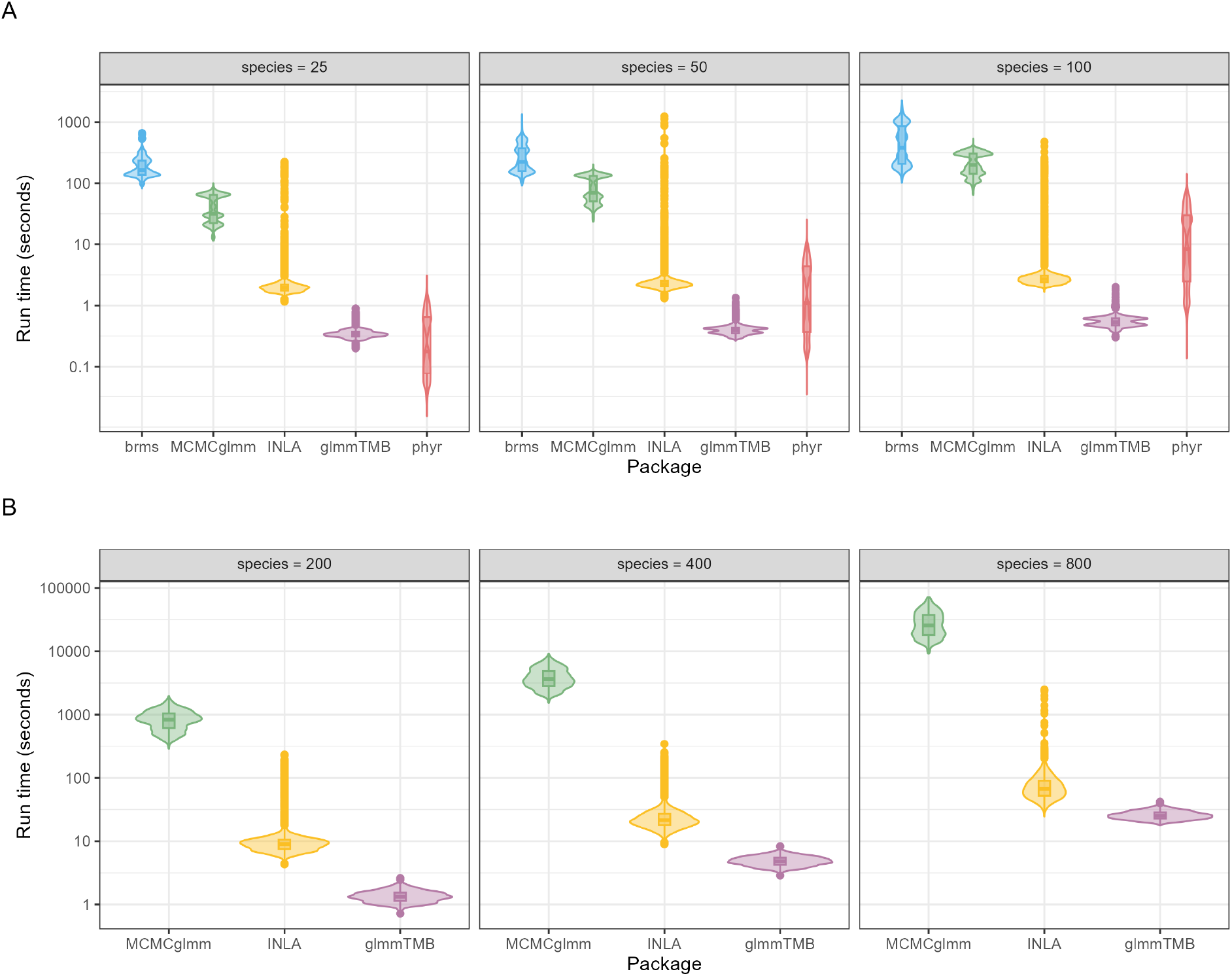
Run times (seconds) for phylogenetic generalised linear mixed models fitted using five R packages (`brms`, `MCMCglmm`, `INLA`, `glmmTMB`, `phyr`). The run time axis is displayed on a log_10_ scale for visualisation. Panels show (A) results from ∼ 50, 000 converged simulated models across datasets with 25 to 100 species, and (B) results from ∼ 6, 000 converged simulated models with 200 to 800 species. All models used the same fixed and random effects structure but different parameter values.

The fixed effect estimates 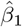 were similar across the five packages, with comparable root mean square error (RMSE) (Figure S2.A and Figure S3.A). Inference was also concordant, as confidence interval widths for 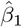 (Figure S2.B and Figure S3.B) and coverage patterns were similar across packages, with all models reaching close to the nominal 95% coverage (Figure S2.C and Figure S3.C). Across all simulated conditions, Figure 2 shows RMSE for the species-level variance 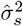 was low for all packages except for `INLA`, which showed higher bias and variance, RMSE for the phylogenetic variance 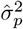 was slightly higher across all packages, with `glmmTMB` and `phyr` exhibiting the lowest median RMSE by a small margin, and RMSE for the residual variance 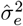 was low across all packages. Figure S4 shows variance component estimates from one simulation scenario, with estimates centred generally on the true values and distinct outliers for `INLA`.

**Figure 2:**
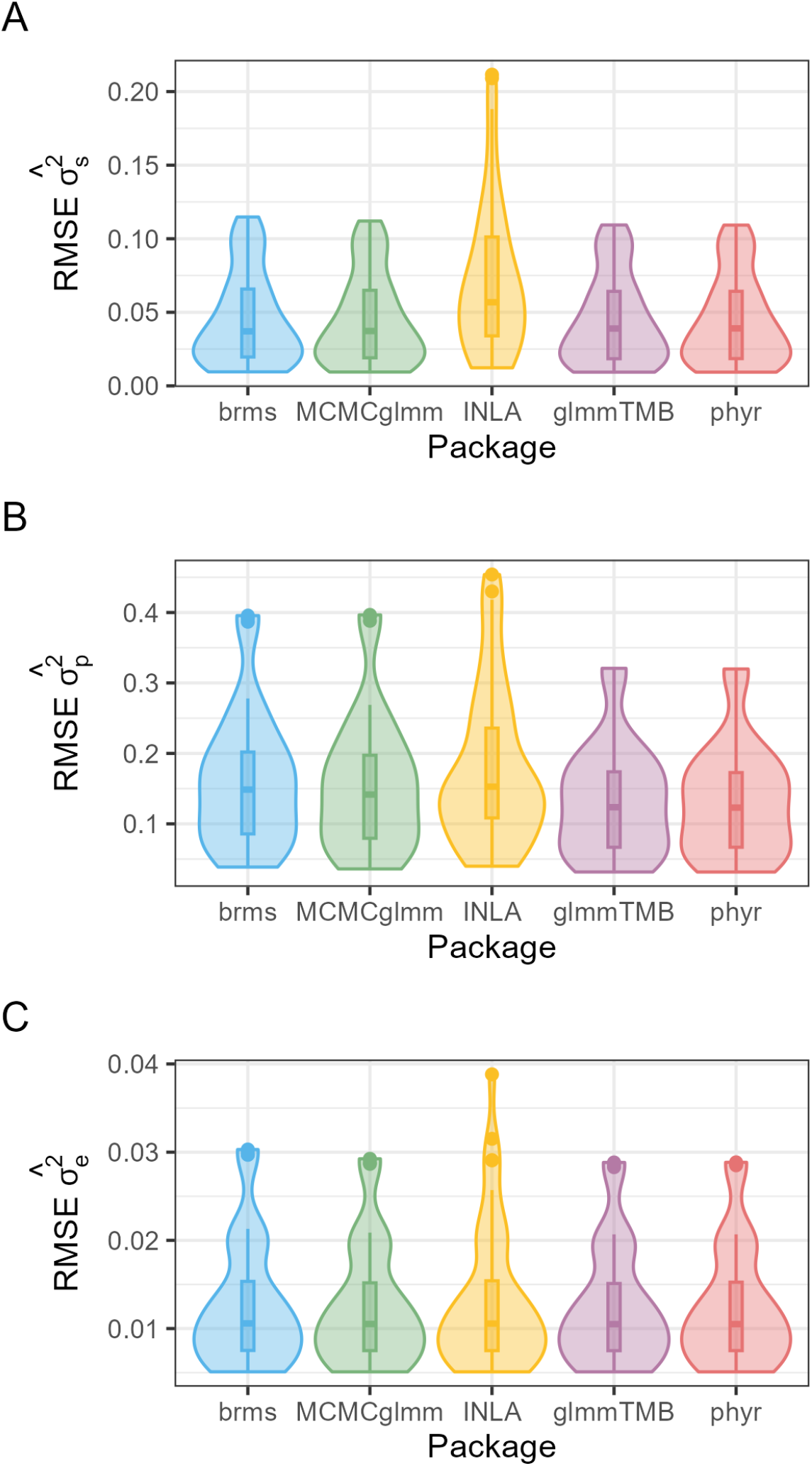
Root mean-squared error (RMSE) of variance component estimates across five R packages. Panels show measures of RMSE of (A) species-level variance 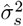, (B) phylogenetic variance 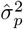, and (C) residual variance 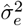. Violin plots summarise simulation results across conditions of 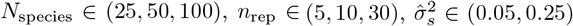, and 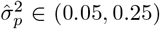 (3 × 3 × 2 × 2 = 36 conditions in total), each based on simulation replicates from converged models (see Table S3).

### 3.2 Applications

To demonstrate the application of the models described above, we use `glmmTMB` to fit PGLMMs with example case studies. The models and results are for illustration only and should not be interpreted or used to make substantive conclusions. We provide all the code and results of the case studies at the following website.

We use avian colouration data to explore trait evolution among birds (Dunn et al., 2015). Birds are a useful system for studying ecological and evolutionary processes because their diversity in morphological, behavioural, and life history traits is well documented across many species and habitats. Avian colouration plays a central role in communication, mate choice and camouflage, and provides valuable insights into the ecological and evolutionary processes that shape bird diversity (Hill & McGraw, 2006). We analyse a dataset of 949 bird species from 446 genera, with spectral reflectance measured for six body regions (crown, back, tail, throat, belly, and wing coverts) from three male and three female individuals, resulting in 30 observations per species (Figure 3.A; Dunn et al., 2015). Spectral data were processed using the pavo R package (Maia et al., 2019) to obtain spectrum shape descriptors.

**Figure 3:**
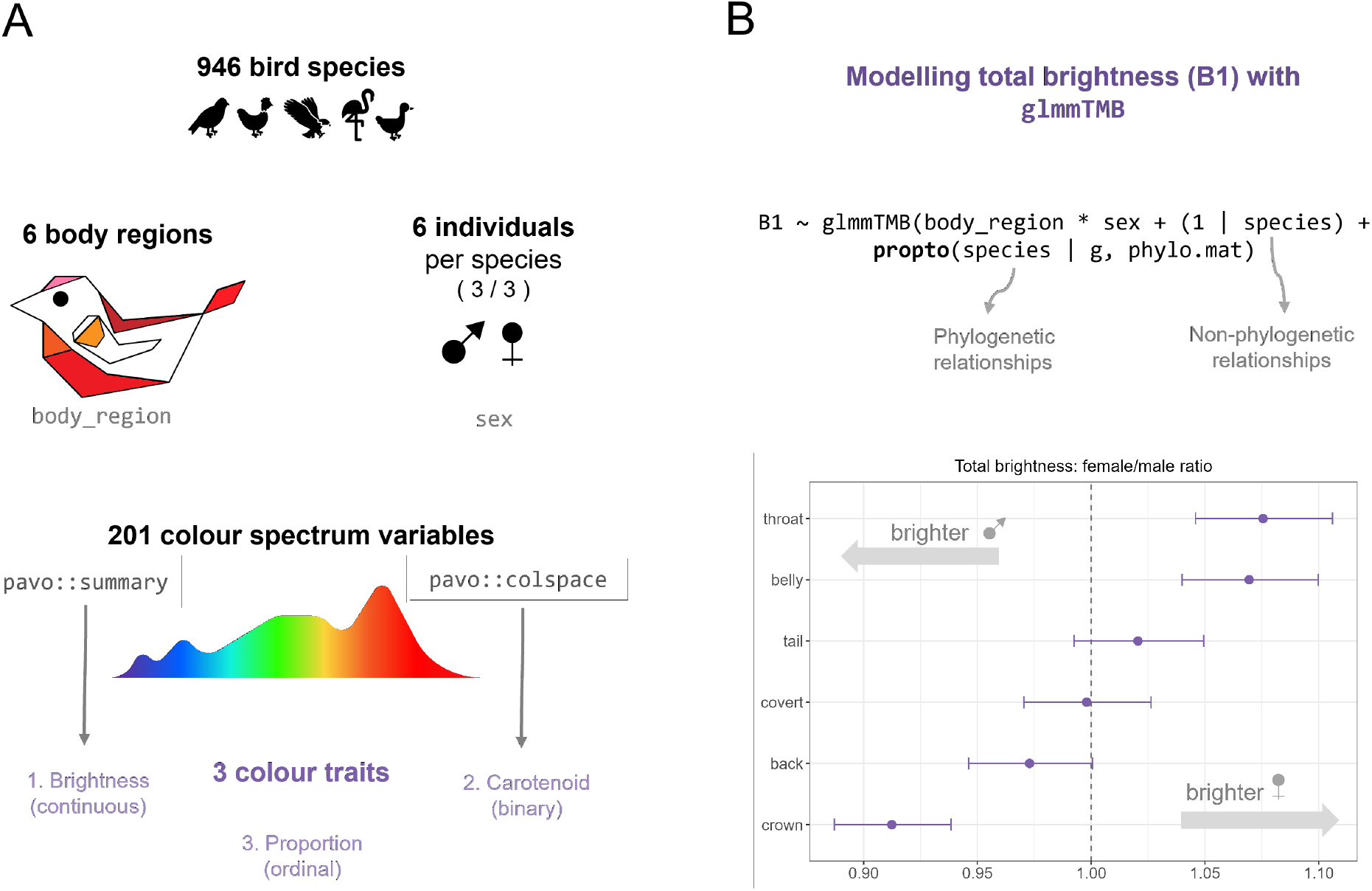
Overview of data and model results for continuous colour trait total brightness. (**A**) Schematic representation of the dataset comprising 946 bird species, with six individuals per species (three males and three females) and spectral measurements taken from six body regions. Each spectrum was described by 201 variables, summarised into three colour traits: a continuous trait (brightness, B1), a binary trait (carotenoid presence), and an ordinal trait (proportion of carotenoid presence across body regions). (**B**) Female-to-male brightness (B1) ratios estimated using a Gamma GLMM with distribution Γ(*μ, ϕ*) and log link log(*μ*) = *η* for B1 across the six body regions. Points represent marginal means and error bars the 95% credible intervals. Ratios above one indicate higher female total brightness, whereas ratios below one indicate higher male total brightness.

#### 3.2.1 Continuous colour trait

We examined the continuous trait of achromatic brightness (B1_ijk_) for observation *i*, measured in body region *j* of species *k* and according to sex. The response B1 is derived from spectral reflectance (300–700 nm) data and represents the total reflectance across the measured spectrum, sometimes called total brightness or spectral intensity (Andersson, 1999; Andersson et al., 1998; Saks et al., 2003). We used a Gamma model with a log link to model the total reflectance across six body regions, following the structure in Equation 3:

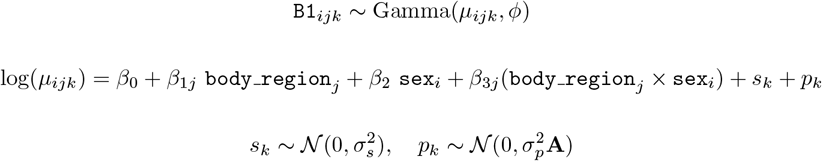

where *β*_0_ is the intercept, *β*_1*j*_ and *β*_3*j*_ represent the effects for each body region *j* (relative to the reference level), *β*_2_ is the main effect of sex, *μ*_*ijk*_ is the mean of the Gamma distribution for observation *i* in body region *j* of species *k, ϕ* is the shape (precision) parameter, *s*_*k*_ is the non-phylogenetic random intercept and *p*_*k*_ is the phylogenetic random intercept. For simplicity in this case study, we assumed the species effect was constant across body regions (noting this may not be biologically realistic in practice). In `glmmTMB` this model can be fitted as:

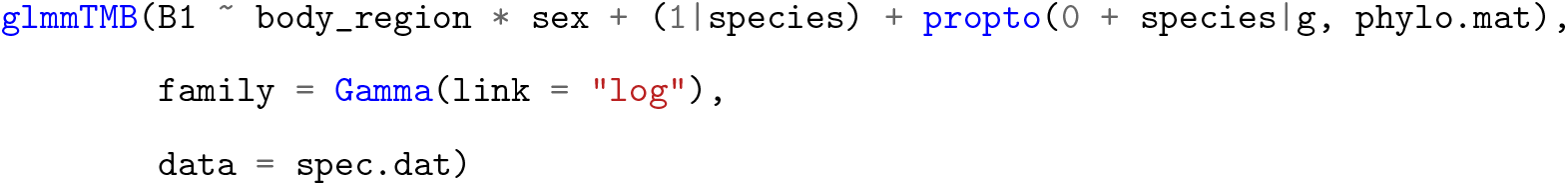

We also investigated a Gaussian model using *log*(*B*1), which resulted in a poorer model fit based on diagnostic residual plots. The phylogenetic species effect explained 84.2% of the total species-level variance, compared to 15.8% for the non-phylogenetic effect. Further, the model found that males had significantly higher total brightness than females for the crown (ratio female/male = 0.91, 95% CI [0.89, 0.94]), while females were significantly brighter in total intensity for the throat (1.08, 95% CI [1.05, 1.11]) and the belly (1.07, 95% CI [1.04, 1.10]) compared to the males (Figure 3.B).

#### 3.2.2 Binary colour trait

Some bird species use carotenoids to colour their plumage from yellow to red, but since they cannot synthesise these pigments, they must acquire them through their diet (Delhey et al., 2023). Carotenoid-based colouration in birds has attracted much attention as a signalling trait and potential bioindicator because its expression depends on environmental availability (Janas et al., 2024; Olson & Owens, 1998; Simons et al., 2014). Understanding how the presence of carotenoid colour varies across species and how it can potentially be shaped by phylogeny provides insight into the evolution of sexually selected signals. Here we model the binary trait carotenoid_*ijk*_ for each `body region`_*j*_ and sex_*i*_ using a binomial GLMM with a logit link, where carotenoid_*ijk*_ indicates the presence (1) or absence (0) of carotenoid colour for observation *i* in body region *j* of species *k*.

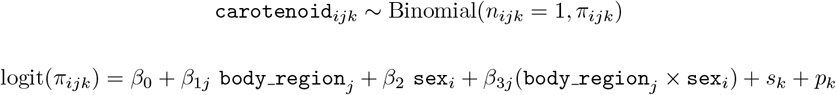

where *π*_*i*_ is the probability of carotenoid presence, *β*_0_ is the intercept, *β*_1*j*_, *β*_2_, and *β*_3*j*_ are the fixed effect coefficients. The model also includes a species-level random intercept (*s*_*k*_) and a phylogenetic random effect (*p*_*k*_) to account for shared evolutionary history, given *k* species and **A** is the phylogenetic variance–covariance matrix. In `glmmTMB` this model can be fitted as:

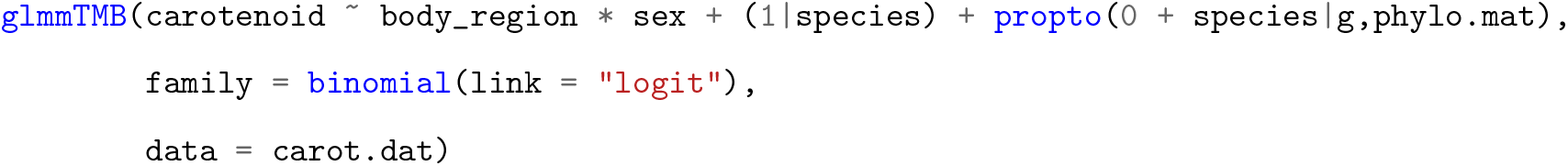

The model showed a strong phylogenetic signal, with 92.4% of the variance from species effect explained by shared evolutionary ancestry. The sex by body region interaction showed no clear trends (see Figure S8). Estimated marginal means indicated odds ratios (female/male) ranging from 0.45 (95% CI [0.18, 1.06]) for the wing coverts to 0.94 (95% CI [0.70, 1.25]) for the belly. The wide confidence intervals around all estimates suggest little evidence of a difference in carotenoid presence between females and males across body regions (Figure S10).

We provide another example from this dataset using the proportion of body regions with carotenoid presence as well as further examples using a plant hydraulic dataset (Sanchez-Martinez et al., 2020) in the Supporting Information section 2.3.

## 4 Discussion

This paper showcases the new covariance `propto` structure (McGillycuddy, 2023) in `glmmTMB` package to fit phylogenetic GLMMs. Our simulation study found that it provides accurate and fast inference. Below we discuss the simulation results, the capabilities of `glmmTMB` to fit flexible distributional families, current limitations, related software beyond the ones assessed in this study, and further considerations for PGLMMs.

### 4.1 Capabilities of glmmTMB for comparative analysis

Our simulation results showed certain models (<2%) fitted with `INLA` package produced unusual or extreme variance estimates for the random components. For `INLA`, this behaviour could potentially be avoided by centring the random effect estimates (by specifying `const = TRUE`) or trying larger starting values (with values *>*4 in: `hyper = list(prec=list(initial=8))`). It is advisable to rerun a model with extreme or unreasonable variance estimates with different packages and such settings to corroborate results. Estimating variance components for random effects is particularly challenging because these parameters are often unstable and can be weakly identified by the likelihood, as it can lie near the boundary at zero and can be sensitive to: small sample sizes; sparse grouping factors; misspecified covariance structure; and the scaling of predictors.

Bayesian frameworks had lower convergence rates (Table 1), reflecting the need for additional tuning of priors and sampling parameters, which in turn increases the time required to achieve stable estimates compared with frequentist approaches. The frequentist package `glmmTMB` avoids the computational cost of MCMC frameworks while delivering similar flexibility in handling multiple distributional families. Traits in ecology and evolution, such as behaviour, life history, and morphology, are known to often depart from the standard Gaussian distribution (e.g. discrete data, or right skewed data). By modelling these traits with suitable distributions and link functions, it reduces the need for data transformation and produces models that more accurately reflect the underlying biological processes. For example, most PGLMM packages do not support a Gamma distribution, which we used in our case study with `glmmTMB` to model achromatic brightness. The only other package that supports this distribution is `brms`, although it is slower to fit than `glmmTMB`.

Further, the `glmmTMB` package accommodates complex models, which current Bayesian packages are unable to fit. For example: reduced-rank approaches (McGillycuddy et al., 2025) for high-dimensional datasets and specialised distributional families such as the Conway–Maxwell–Poisson distribution for both under- and over-dispersed count data (Brooks et al., 2019). Ongoing community contributions continue to expand the functionalities of `glmmTMB`, adding new model structures and distributional families. Further, this package retains a flexible range of options with an easy-to-use and familiar interface similar to `lme4` (Bates et al., 2015). Table 2 summarises functionality across the packages assessed in our simulation study to fit PGLMMs for comparative analysis, showing that `glmmTMB` supports wide distributional families, flexible random-effect structures, and fast ML/REML estimation, while Bayesian frameworks provide similar scope at a higher computational cost.

**Table 2:** Comparison of R packages functionalities for fitting phylogenetic generalised linear mixed models (PGLMMs). Star (*) ratings indicate relative functionality or performance, with more stars representing greater flexibility, broader support, or faster speed. **RE structures** refer to the random effect structures available, one star refers to the following structures: spatial, AR1 (temporal), and phylogenetic. **Smooth terms** are functional terms used to model non-linear relationships, often implemented with spline functions. **Speed** refers to the overall computational time, which included compilation and sampling for MCMC models, these may differ depending on model specification especially for Bayesian frameworks (e.g. expect slower time with higher number of iterations).

| Package | Estimation | RE structures | Smooth terms | Supported families * | Speed |
| --- | --- | --- | --- | --- | --- |
| glmmTMB | MLE/REML/MCMC<sup>†</sup> | *** | Yes | *** | *** |
| brms | MCMC | *** | Yes | *** | * |
| MCMCglmm | MCMC | ** | Yes | ** | ** |
| INLA | INLA | *** | Yes | *** | *** |
| phyr | MLE/REML/INLA<sup>‡</sup> | * | No | * | *** |

\*Table S1 provides details of supported distributional families per package.
<sup>†</sup>glmmTMB can fit models using a Bayesian approach with a Hamiltonian Monte Carlo (HMC) sampler.
<sup>‡</sup>phyr has the functionality to fit models using a Bayesian approach with an INLA integration.

### 4.2 Limitations and other relevant software

Our simulation study is limited by the particular parameter conditions and simulated scenarios assessed. Datasets with more than 800 species will run slower. In the Bayesian framework, we applied the same priors and parameter settings across all simulated scenarios, which may explain the higher non-convergence rates for `MCMCglmm` at larger sample sizes. We did not test other software that can also fit PGLMMs. ASReml-R (Butler et al., 2023), for example, uses a penalized quasi likelihood (PQL) approach and is computationally very fast, likely faster than `glmmTMB`, and has been widely used in genetics, agriculture, and comparative analyses. However, it is closed source and requires a paid licence (sold by VSN International). We also did not investigate R packages developed primarily for community phylogenetic ecology and joint species distribution modelling with large multivariate datasets, such as the `pez` package (Ives & Helmus, 2011; Rafferty & Ives, 2013), the HMSC package for hierarchical modelling of species communities using MCMC sampling (Tikhonov et al., 2020), and gllvm for Generalized Linear Latent Variable Models using variational approximations (Niku et al., 2019; van der Veen & O’Hara, 2024). These tools are particularly suited for questions of species co-occurrence and ecological assembly processes while controlling for phylogeny. PGLMMs are also able to address such questions (Gallinat & Pearse, 2021), and `glmmTMB` can fit these types of models through its reduced rank covariance structure (McGillycuddy et al., 2025). However, our study did not cover this area and instead focused on comparative trait analysis. Other R packages with PGLMM capability, not assessed here but worth mentioning, are phylopath (Bijl, 2018) and the general purpose GLMM function `glmmPQL` in the `MASS` package (Venables & Ripley, 2002) in conjunction with `corBrownian` function from the `ape` package (Paradis & Schliep, 2019). Further, we note that the current `propto` implementation in `glmmTMB` can accommodate independent random slopes for continuous variables, but does not yet support random slopes for categorical predictors and correlations between random slopes and intercepts, which is an area of future development. Another formulation of PGLMMs for evolutionary analyses implemented with `glmmTMB` (Li & Bolker, 2019) can fit PGLMMs with correlated random slope and intercepts. This implementation was not evaluated in our study.

The use of PGLMMs in comparative analyses provides a flexible framework for modelling phylogenetic structure, but they rest on several strong assumptions. One assumption is that the phylogenetic relationships in a given tree are correct, yet in practice phylogenies are always estimated with uncertainty, and different plausible trees may yield different inferences. Bayesian methods can accommodate this uncertainty by integrating over multiple candidate phylogenetic trees within the posterior, providing topology-corrected estimates. In frequentist frameworks, this uncertainty can be approximated by averaging model estimates across sets of trees using approaches such as Rubin’s rule (Nakagawa & De Villemereuil, 2019). However, regardless of the approach, the true tree is never known with certainty. A second assumption is that the evolutionary model underlying trait variation is correctly specified. While Brownian motion is often assumed in evolutionary ecology, alternative models such as Ornstein–Uhlenbeck or early-burst may be more appropriate in some cases (Blomberg et al., 2020; Boettiger et al., 2012; Pennell et al., 2015), depending on the question. PGLMMs can in principle accommodate alternative covariance structures, but testing and fitting these models is non-trivial and often requires generating covariance matrices from specialised packages such as `geiger` (Pennell et al., 2014) or `mvMORPH` (Clavel et al., 2015). These assumptions highlight the importance of interpreting PGLMM results with caution, as both tree misspecification and model misspecification can influence estimates and conclusions of the phylogenetic signal and trait evolution.

## 5 Conclusion

Modelling phylogenetic relationships is important because failure to account for shared evolutionary history among species can lead to inflated type I error rates arising from incorrect variance estimation (Martins & Garland Jr., 1991; Rohle, 2006). Our study demonstrates that `glmmTMB` can flexibly fit PGLMMs using the `propto` (proportional) covariance structure, with results comparable to those from Bayesian packages, but with substantially faster computation. This speed is particularly advantageous for analyses that require repeated model fitting (e.g. simulations or resampling), which are often prohibitively slow with Bayesian frameworks. Further, `glmmTMB` supports advanced phylogenetic analyses currently not implemented in other maximum-likelihood frameworks, such as reduced-rank approaches (McGillycuddy et al., 2025) and location-scale (mean-dispersion) modelling (Nakagawa et al., 2025). In future work, there is considerable scope for advancing PGLMM implementations, particularly in extending phylogenetic mixed models to more complex frameworks, such as those accommodating multi response structures with different distributional families. The `glmmTMB` package is a general-purpose GLMM framework that is well-suited to model the complex data typical of ecology and evolution.

## Box 1: Estimation of PGLMMs

Estimating phylogenetic generalised linear mixed models (PGLMMs) is challenging for two reasons. First, the marginal likelihood involves integrating over high-dimensional random effects and has no general closed-form solution (Pinheiro & Bates, 2006). Second, evaluating the integral is computationally costly as it must be approximated numerically. Frequentist tools uses deterministic approximations, such as the Laplace method or adaptive Gaussian quadrature for example (see more details in Bolker et al., 2009), which are fast but sometimes inaccurate. Bayesian approaches avoid the need for analytic integration by sampling from the posterior with Markov chain Monte Carlo (MCMC) or related algorithms, which can yield more accurate estimates but requires much longer computational time. Below we describe the estimation approach of `glmmTMB`, along with the four packages assessed in this paper to fit PGLMMs.

### 1. Frequentist framework

`glmmTMB` estimates model parameters using maximum likelihood (ML) or restricted maximum likelihood (REML), with the Laplace approximation used to marginalise over random effects (Brooks et al., 2017). It is built on the Template Model Builder (TMB) framework for efficient optimisation, using automatic differentiation (Kristensen et al., 2016). `glmmTMB` supports a wide range of distributions, custom covariance structures (Kristensen & McGillycuddy, 2025), and efficient sparse matrix computation. The package is particularly well-suited for complex datasets (i.e. multiple random effects) and has been shown to be faster than other GLMM packages such as glmer. With the `phyr` package, PGLMMs are estimated using a maximum likelihood implementation via C++ with restricted maximum likelihood (REML) and incorporates phylogenetic correlation structures (Ives et al., 2020; Li et al., 2020). `phyr` has a user-friendly interface similar to the lme4 package (Bates et al., 2015). While `phyr` supports binary, binomial, and Poisson models, it uses a Bayesian approach to run zero-inflated binomial and Poisson models via the integrated nested Laplace approximation (`INLA`; see below). offering an alternative estimation framework within the same syntax. The `phyr` package is limited to specific model structures and cannot fit a phylogenetic random effect alone (it must be paired with a species-level random effect).

### 2. Bayesian framework

`MCMCglmm` uses Gibbs sampling to generate posterior draws by sequentially sampling from conditional distributions. This package can accommodate phylogenetic covariance structure as a random effect, residual variance structures, and multivariate models. The package is constructed to run only a single chain, with the user needing to adjust the number of iterations for convergence, although this can be worked around by running multiple independent chains sequentially. Priors are limited, with inverse-Wishart priors for (co)variance components and for residual variances, and Gaussian priors for fixed effects. `MCMCglmm` has been shown to be fast (Hadfield, 2015; Hadfield & Nakagawa, 2010). The `brms` package fits models an augmented Hamiltonian Monte Carlo (HMC) with the No-U-Turn Sampler, an algorithm to improve sampling efficiency, implemented via Stan (Carpenter et al., 2017). This allows efficient exploration of high-dimensional posteriors and supports automatic computation of convergence diagnostics across multiple chains. `brms` accommodates a wide range of distributions, link functions, and user-defined priors, but model fitting is resource-intensive, particularly for large phylogenetic trees or complex hierarchical models, which can slow down model run time. To avoid sampling we can use the deterministic approximations based on `INLA`, the Integrated Nested Laplace Approximation (Rue et al., 2009). `INLA` approximates posterior marginals for latent Gaussian models by combining nested Laplace approximations and numerical integration. It offers rapid computation and is particularly effective for spatial or hierarchical models with large datasets (Rue et al., 2017).

