## Supporting information for "Fast phylogenetic generalised linear mixed-effects modelling using the glmmTMB R package"

### Supporting Information to: "*Phylogenetic generalised linear mixed-effects modelling with glmmTMB R package*"

### 1 Supporting methods

#### 1.1 R package distributional families

Below is the table of distributional families across the five R packages assessed in our simulation study.

Table S1: Current supported distributional families from the five packages evaluated in our simulation study. The following distribution families are not exhaustive and represent the options currently available in each package, which may expand or change in future releases.

| | Packages | | | | |
|---|---|---|---|---|---|
| **Distribution family** | `glmmTMB` | `phyr` | `brms` | `MCMCglmm` | `INLA` |
| Gaussian (normal) | × | × | × | × | × |
| Poisson | × | × | × | × | × |
| Binomial | × | × | × | × | × |
| Zero-inflated Poisson | × | ×* | × | × | × |
| Zero-inflated Binomial | × | ×* | × | × | × |
| Negative binomial | × | | × | × | × |
| Hurdle models | × | | × | × | × |
| Gamma | × | | × | | × |
| Beta | × | | × | | × |
| Beta-binomial | × | | × | | × |
| Tweedie | × | | × | | × |
| Conway–Maxwell–Poisson | × | | | | |

* only with the `phyr` Bayesian approach.

#### 1.2 Simulation study

##### Bayesian prior settings

For all `brms` models in the simulation study, we used the default weakly informative priors, which provide mild regularisation by stabilising estimation and preventing extreme parameter values while remaining flexible across models.

For `MCMCglmm`, we used the recommended parameter-expanded priors, which introduce auxiliary parameters to improve mixing and convergence in models with multiple random effects, for the model in equation 3 the priors are specified as:

```
prior <- list(
  G = list(
    G1 = list(V = 1, nu = 1, alpha.mu = 0, alpha.V = 1000),
    G2 = list(V = 1, nu = 1, alpha.mu = 0, alpha.V = 1000)
  ),
  R = list(V = 1, nu = 0.02)
)
```

where $V = 1$ specifies the scale matrix, and $\nu = 1$ gives weakly informative inverse-Wishart-type priors for the random-effect variances. The terms $\alpha_\mu = 0$ and $\alpha_V = 1000$ define the auxiliary parameters with a Normal$(0, 1000)$ prior to improve sampling efficiency. For the residual variance ($R$), we used a very weak prior ($V = 1, \nu = 0.02$), which exerts minimal influence and allows the data to dominate estimation. For models with one measure per species (equation 2) we used the same prior without the second random effect `G2`.

We note that in practice the selection of priors should reflect relevant prior knowledge where available, as their influence on the model estimates depends on the specific context; a prior that is diffuse in one analysis may be informative in another. The effect of prior choice can be assessed using prior predictive checks (Gelman et al., 1996; Wesner & Pomeranz, 2021).

##### Bayesian model specification

The settings were chosen for simplicity, with the number of iterations determined from pilot runs to ensure that at least 80% of simulations achieved an effective sample size (ESS) greater than 400 for the main parameters. We note that this threshold is arbitrary and would require adjustment depending on factors such as sample size, model complexity, the number and structure of random effects, the strength of priors, and the degree of autocorrelation in the chains.

Each model for `brms` was fitted using 10000 warmup iterations (`warmup = 10000`) and 10000 post-warmup iterations (`iter = 20000`) in four chains (`chains = 4`), adding up to 40000 samples from the posterior distribution, with computation run on four cores (`cores = 4`).

Each model for `MCMCglmm` was fitted using a total of 303000 iterations (`nitt = 303000`), with a burn-in of 3000 iterations (`burnin = 3000`) and a thinning interval of 10 (`thin = 10`), resulting in 30000 posterior samples per chain.

##### Convergence metrics

Convergence metrics were set for each package respectively on the following criteria. The `phyr` package, we only assessed either the model return without any error or warning message. The convergence metric for the `glmmTMB` package assesses whether the Hessian is positive definite, which suggests that the maximum likelihood estimation has likely converged to a local minimum (ideally a global one), and that standard errors are reliably estimated, and we also flagged any errors or warnings from the output.

```
conv_tmb <- if (!is.null(model_glmmTMB)) model_glmmTMB$sdr$pdHess else FALSE
```

Convergence was assessed for models from the `brms` package using the maximum R-hat statistic across all parameters, with values below 1.01 indicating successful convergence (Vehtari et al., 2021). R-hat values close to 1.00 suggest that chains have mixed well and sampled from the same posterior distribution, whereas values above 1.01 indicate potential problems with convergence or model fit.

```
conv_brms <- if (!is.null(model_brm)) max(rhat(model_brm)) < 1.01 else FALSE
```

The `MCMCglmm` package was assessed for convergence using the Heidelberger and Welch diagnostic applied to the combined posterior samples of the fixed effects (`Sol`) and variance components (`VCV`). Convergence was deemed to be achieved only if the diagnostic indicated stationarity for all parameters in the chain.

```
conv_mcmc <- if (!is.null(model_mcmc)) {
  fullchain <- cbind(as.mcmc(model_mcmc$Sol), as.mcmc(model_mcmc$VCV))
  all(heidel.diag(fullchain)[,1] == 1)
} else FALSE
```

For both the `brms` and `MCMCglmm` models were retained only if the effective sample size (ESS) for all model parameters exceeded 400.

The `INLA` package was assessed for convergence using the internal convergence flag returned in the model object (`conv_inla <- model_inla$ok`). Any warning or error messages were also recorded and assessed for non convergence. In addition, we observed extreme variance component estimates for some models that passed the convergence test. Therefore, models were flagged as non converged if any random effect variance component exceeded a variance of 4 ($\sigma_p^2, \sigma_s^2 > 4$), which is more than 16 times the true variance value of the conditions set in our simulation study.

Table S2: Simulation parameters used to compare model performance across packages. "Conditions" indicates the number of unique simulated parameter conditions per package and is calculated as the product of all combinations of these varying factors. "unbal." refers to an unbalanced design for the number of replicates per species (see Methods).

| **Package** | **Model specification** | $N_{\mathbf{sim}}$ | **Conditions** | $n_{\mathbf{reps}}$ | $N_{\mathbf{species}}$ | $\beta_0$ | $\beta_1$ | $\sigma_s^2$ | $\sigma_p^2$ | $\sigma_e^2$ |
|---|---|---|---|---|---|---|---|---|---|---|
| `phyr`, `glmmTMB`, `brms`, `MCMCglmm`, `INLA` | repeated measures (Eq. 3) | 4000 | 48 | 5, 10, 30, unbal. | 25, 50, 100 | 1 | 1.5 | 0.05, 0.25 | 0.05, 0.25 | 0.2 |
| `glmmTMB`, `MCMCglmm`, `INLA` | repeated measures (Eq. 3) | 500 | 48 | 5, 10, 30, unbal. | 200, 400, 800 | 1 | 1.5 | 0.05, 0.25 | 0.05, 0.25 | 0.2 |
| `glmmTMB`, `brms`, `MCMCglmm`, `INLA` | one measure (Eq. 2) | 4000 | 6 | 1 | 25, 50, 100 | 1 | 1.5 | - | 0.05, 0.25 | 0.2 |
| `glmmTMB`, `MCMCglmm`, `INLA` | one measure (Eq. 2) | 500 | 6 | 1 | 200, 400, 800 | 1 | 1.5 | - | 0.05, 0.25 | 0.2 |

### 2 Supporting results

#### 2.1 Simulation study: repeated measures per species

The following section describes supporting results of the simulations of models with repeated measurements per species ($n_{\text{reps}} > 1$), described in Equation 3 in the main text.

##### 2.1.1 Model convergence

Table S3: Number of retained simulated models per species size per package after removing any runs where at least one of the five packages failed to converge.

| | $N_{\text{species}}$ | | | | | |
|---|---|---|---|---|---|---|
| **Simulation per package** | **25** | **50** | **100** | **200** | **400** | **800** |
| Count | 44,566 | 48,557 | 51,689 | 6,890 | 6,219 | 5,389 |
| Proportion retained (%) | 69.6% | 75.9% | 80.8% | 86.1% | 77.7% | 67.4% |

##### 2.1.2 Model run time

Table S4: Mean model run time for each R package across increasing number of species ($N_{\text{species}}$). Values are expressed in seconds (s), minutes (m), or hours (h), depending on magnitude.

| | $N_{\text{species}}$ | | | | | |
|---|---|---|---|---|---|---|
| **Package** | **25** | **50** | **100** | **200** | **400** | **800** |
| `brms` | 3m 09s | 4m 39s | 8m 55s | — | — | — |
| `MCMCglmm` | 42.12s | 1m 27s | 3m 42s | 14m 11s | 1h 05m | 7h 59m |
| `INLA` | 2.30s | 2.76s | 3.37s | 12.07s | 26.57s | 1m 20s |
| `glmmTMB` | 0.35s | 0.40s | 0.55s | 1.36s | 4.90s | 25.92s |
| `phyr` | 0.39s | 2.61s | 18.44s | — | — | — |

For one example simulated dataset of $N_{species} = 30$, we assessed the breakdown of the two Bayesian packages run times by separating the compilation and sampling times. We found for `MCMCglmm` compilation ran in 0.06 seconds (0.3%) and sampling ran in 23.59 seconds (99.7%). For `brms` compilation ran in 168.14 seconds (13.9%) and sampling ran 869.25 seconds (86.1%). Code for separating compilation and run time is provided in supporting webpage: https://anonymous.4open.science/w/phylo_glmm_sim_mee/supp_information.html#runtimes.

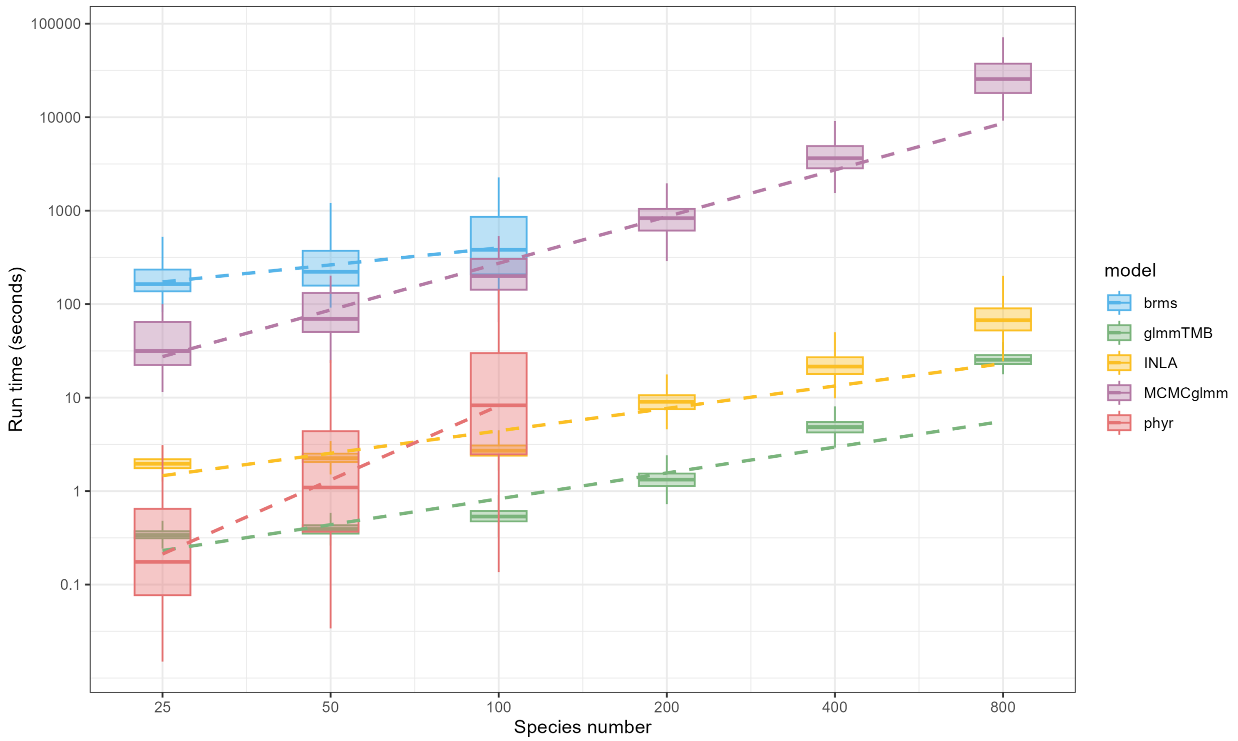

Figure S1: Computation times (seconds) for fitting phylogenetic models across packages in relation to species number, plotted on logarithmic axes. Boxplots show run time distributions per package, with coloured dashed lines indicating fitted regressions.

##### 2.1.3 Fixed covariate estimate

Table S5: Monte Carlo Standard Errors (MCSE) of the estimate $\hat{\beta}_1$ bias per package and per number of species.

| package | species size | $\hat{\beta}_1$ MCSE |
|---|---|---|
| INLA | 25 | 0.0303 |
| MCMCglmm | 25 | 0.0301 |
| brms | 25 | 0.0301 |
| glmmTMB | 25 | 0.0301 |
| phyr | 25 | 0.0301 |
| INLA | 50 | 0.0212 |
| MCMCglmm | 50 | 0.0211 |
| brms | 50 | 0.0211 |
| glmmTMB | 50 | 0.0211 |
| phyr | 50 | 0.0211 |
| INLA | 100 | 0.0146 |
| MCMCglmm | 100 | 0.0146 |
| brms | 100 | 0.0146 |
| glmmTMB | 100 | 0.0146 |
| phyr | 100 | 0.0146 |
| INLA | 200 | 0.0102 |
| MCMCglmm | 200 | 0.0102 |
| glmmTMB | 200 | 0.0102 |
| INLA | 400 | 0.0076 |
| MCMCglmm | 400 | 0.0076 |
| glmmTMB | 400 | 0.0076 |
| INLA | 800 | 0.0054 |
| MCMCglmm | 800 | 0.0054 |
| glmmTMB | 800 | 0.0054 |

Table S6: Monte Carlo Standard Errors (MCSE) of $\hat{\beta}_1$ coverage rates per package and per number of species.

| package | species size | mean cov $\hat{\mu}$ | cov $\hat{\mu}$ MCSE |
|---|---|---|---|
| brms | 25 | 0.949 | 0.0637 |
| MCMCglmm | 25 | 0.948 | 0.0638 |
| INLA | 25 | 0.947 | 0.0649 |
| glmmTMB | 25 | 0.946 | 0.0650 |
| phyr | 25 | 0.946 | 0.0654 |
| brms | 50 | 0.947 | 0.0559 |
| MCMCglmm | 50 | 0.947 | 0.0561 |
| INLA | 50 | 0.946 | 0.0565 |
| glmmTMB | 50 | 0.946 | 0.0564 |
| phyr | 50 | 0.946 | 0.0565 |
| brms | 100 | 0.952 | 0.0532 |
| MCMCglmm | 100 | 0.952 | 0.0534 |
| INLA | 100 | 0.952 | 0.0535 |
| glmmTMB | 100 | 0.952 | 0.0535 |
| phyr | 100 | 0.952 | 0.0535 |
| MCMCglmm | 200 | 0.953 | 0.0529 |
| INLA | 200 | 0.952 | 0.0532 |
| glmmTMB | 200 | 0.952 | 0.0537 |
| MCMCglmm | 400 | 0.946 | 0.0654 |
| INLA | 400 | 0.947 | 0.0649 |
| glmmTMB | 400 | 0.946 | 0.0650 |
| MCMCglmm | 800 | 0.952 | 0.0535 |
| INLA | 800 | 0.953 | 0.0530 |
| glmmTMB | 800 | 0.953 | 0.0530 |

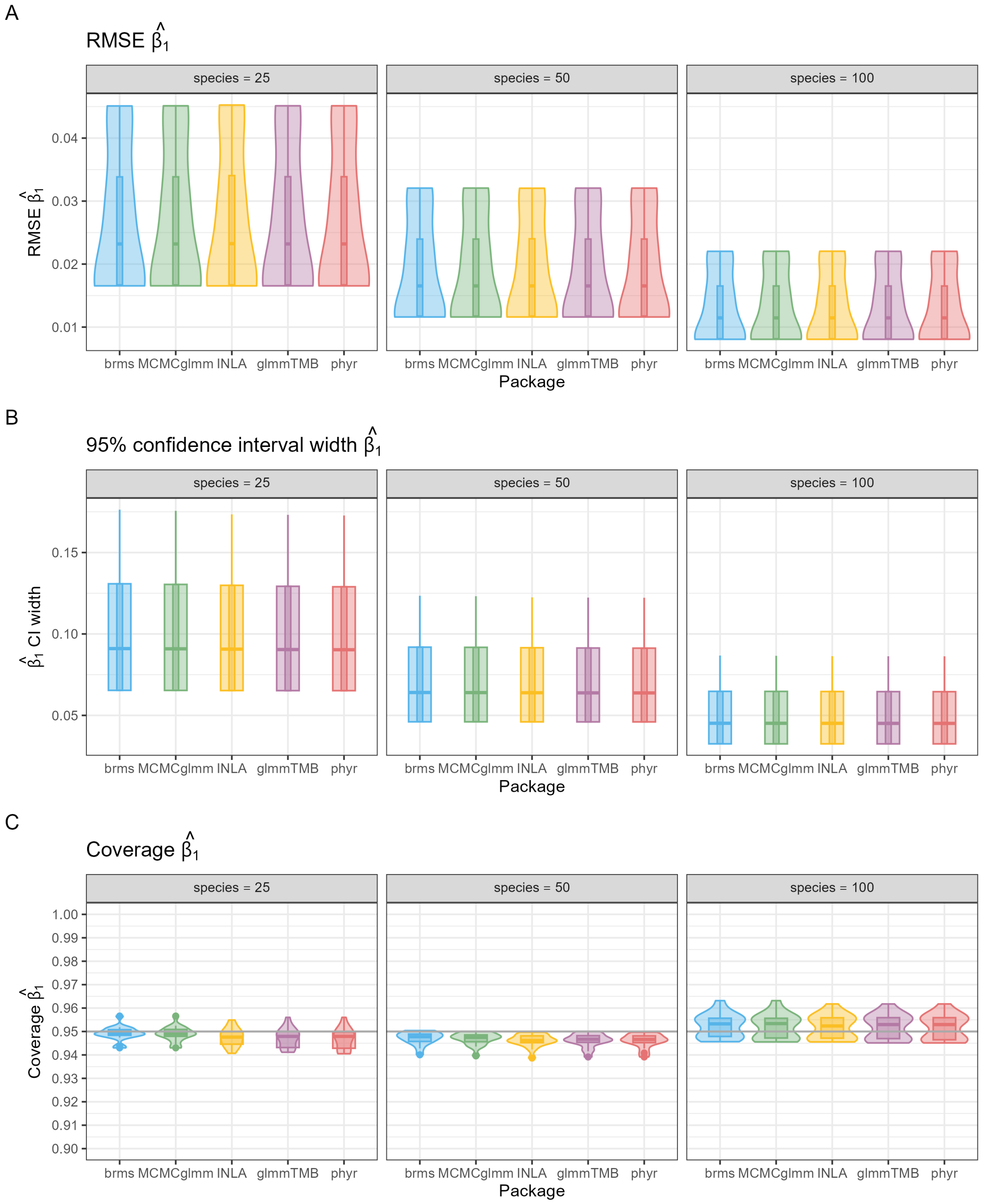

Figure S2: Comparison of five R packages in estimating the fixed effect $\hat{\beta}_1$ for datasets with 25, 50, and 100 species. Panels show (A) root-mean-square error (RMSE), (B) 95% confidence interval width, and (C) 95% interval coverage of the fixed effect estimate $\hat{\beta}_1$ . Violin plots summarise simulation results across three conditions of $N_{\text{species}}$, and all conditions of $n_{\text{rep}}$, $\hat{\sigma}_s^2$, and $\hat{\sigma}_p^2$ (48 conditions in total), each based on simulation replicates of converged models.

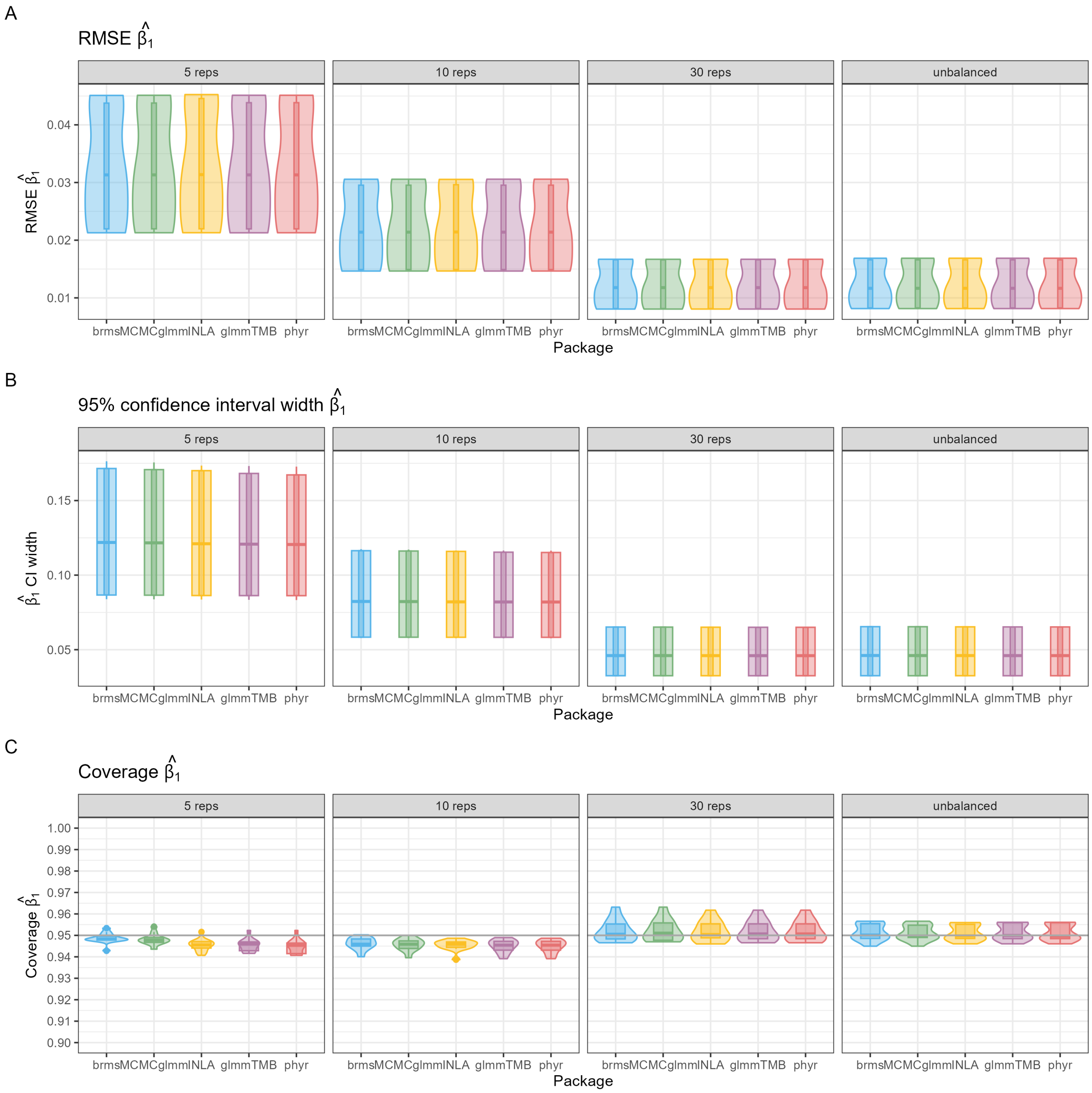

Figure S3: Comparison of five R packages in estimating the fixed effect $\hat{\beta}_1$ for datasets $n_{reps} \in (5, 10, 30)$ and the unbalanced design (see Methods section). Panels show (A) root-mean-square error (RMSE), (B) 95% confidence interval width, and (C) 95% interval coverage of the fixed effect estimate $\hat{\beta}_1$. Violin plots summarise simulation results across three conditions of $N_{\text{species}}$, and all conditions of $n_{\text{rep}}$, $\hat{\sigma}^2_s$, and $\hat{\sigma}^2_p$ (48 conditions in total), each based on simulation replicates of converged models

##### 2.1.4 Variance component estimates

Table S7: Monte Carlo Standard Errors (MCSE) of variance components ($\hat{\sigma}_s^2$, $\hat{\sigma}_p^2$ and $\hat{\sigma}_e^2$) estimates per package and per number of species.

| package | species size | $\hat{\sigma}_s^2$ MCSE | $\hat{\sigma}_p^2$ MCSE | $\hat{\sigma}_e^2$ MCSE |
|---|---|---|---|---|
| brms | 25 | 0.115 | 0.246 | 0.020 |
| MCMCglmm | 25 | 0.111 | 0.250 | 0.019 |
| INLA | 25 | 0.164 | 0.312 | 0.021 |
| glmmTMB | 25 | 0.107 | 0.223 | 0.019 |
| phyr | 25 | 0.107 | 0.222 | 0.019 |
| brms | 50 | 0.108 | 0.197 | 0.014 |
| MCMCglmm | 50 | 0.106 | 0.194 | 0.014 |
| INLA | 50 | 0.129 | 0.218 | 0.014 |
| glmmTMB | 50 | 0.103 | 0.171 | 0.013 |
| phyr | 50 | 0.103 | 0.170 | 0.014 |
| brms | 100 | 0.104 | 0.159 | 0.009 |
| MCMCglmm | 100 | 0.103 | 0.155 | 0.009 |
| INLA | 100 | 0.110 | 0.152 | 0.009 |
| glmmTMB | 100 | 0.102 | 0.142 | 0.009 |
| phyr | 100 | 0.102 | 0.142 | 0.009 |
| MCMCglmm | 200 | 0.103 | 0.131 | 0.007 |
| INLA | 200 | 0.105 | 0.121 | 0.007 |
| glmmTMB | 200 | 0.102 | 0.124 | 0.007 |
| MCMCglmm | 400 | 0.101 | 0.118 | 0.005 |
| INLA | 400 | 0.102 | 0.116 | 0.005 |
| glmmTMB | 400 | 0.101 | 0.113 | 0.005 |
| MCMCglmm | 800 | 0.100 | 0.110 | 0.003 |
| INLA | 800 | 0.101 | 0.107 | 0.003 |
| glmmTMB | 800 | 0.100 | 0.107 | 0.003 |

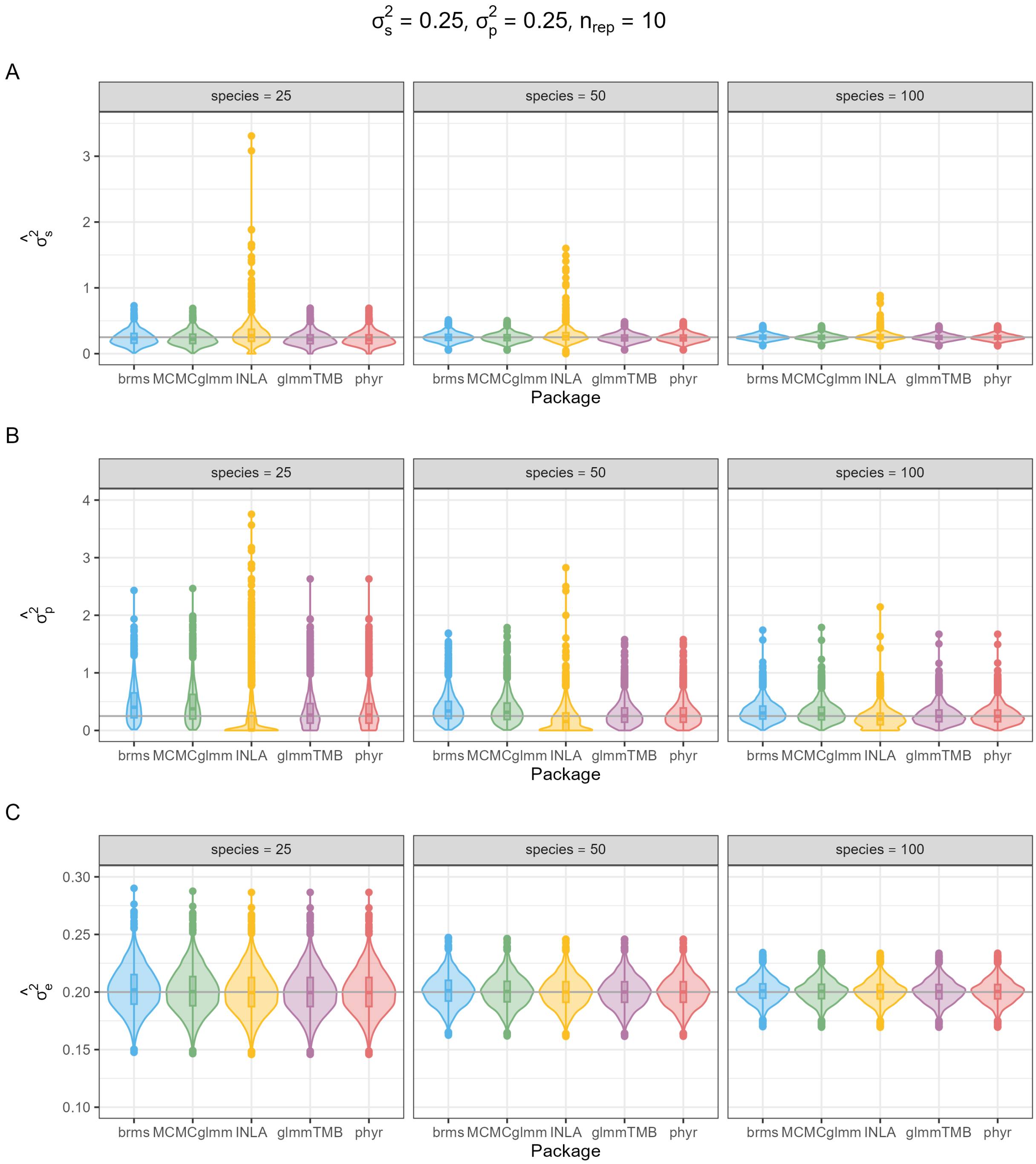

Figure S4: Variance component estimates across five R packages for simulated datasets with 25, 50, and 100 species. Panels show distribution of (A) species-level variance $\hat{\sigma}_s^2$, (B) phylogenetic variance $\hat{\sigma}_p^2$, and (C) residual variance $\hat{\sigma}_e^2$ estimated, for true values of $\sigma_s^2 = 0.25$, $\sigma_p^2 = 0.25$, and $n_{rep} = 10$ replicates per species. Violin plots with boxplots summarise results across approximately ∼4,000 simulations.

#### 2.2 Simulation study: one measure per species

The following section describes supporting results of the simulations of models with with a single measurement per species ($n_{\text{reps}} = 1$), described in Equation 2 in the main text. We did not fit models with `phyr` for datasets containing only one measure per species as we couldn't specify a single phylogenetic random effect (it fits automatically a second species level-effect).

##### 2.2.1 Model convergence

Table S8: Convergence rates percentage (%) of phylogenetic mixed models per package across $N_{\text{species}}$. Results are based on 8,000 simulations for each $N_{\text{species}} \leq 100$ and 1,000 simulations for each $N_{\text{species}} \geq 100$, given varying variance component values ($\sigma_p^2$). Full details of varying parameter values is provided in supporting table S2.

| | $N_{\text{species}}$ | | | | | |
|---|---|---|---|---|---|---|
| **Package** | **25** | **50** | **100** | **200** | **400** | **800** |
| `brms` | 79.6 | 95.9 | 98.9 | – | – | – |
| `MCMCglmm` | 96.9 | 97.2 | 97.1 | 95.1 | 97.2 | 96.1 |
| `INLA` | 100† | 100 | 100 | 100 | 100 | 100 |
| `glmmTMB` | 99.9 | 100 | 100 | 100 | 100 | 100 |

†four models did not converge.

Table S9: Number of retained simulated models per species size per package after removing any runs where at least one of the five packages failed to converge or if either `brms` or `MCMCglmm` had an effective sample size (ESS) below 400.

| | $N_{\text{species}}$ | | | | | |
|---|---|---|---|---|---|---|
| **Simulation per package** | **25** | **50** | **100** | **200** | **400** | **800** |
| Count | 24,628 | 29,832 | 30,724 | 2,853 | 2,916 | 2,883 |
| Proportion retained (%) | 77.0% | 93.2% | 96.0% | 95.1% | 97.2% | 96.1% |

##### 2.2.2 Model run time

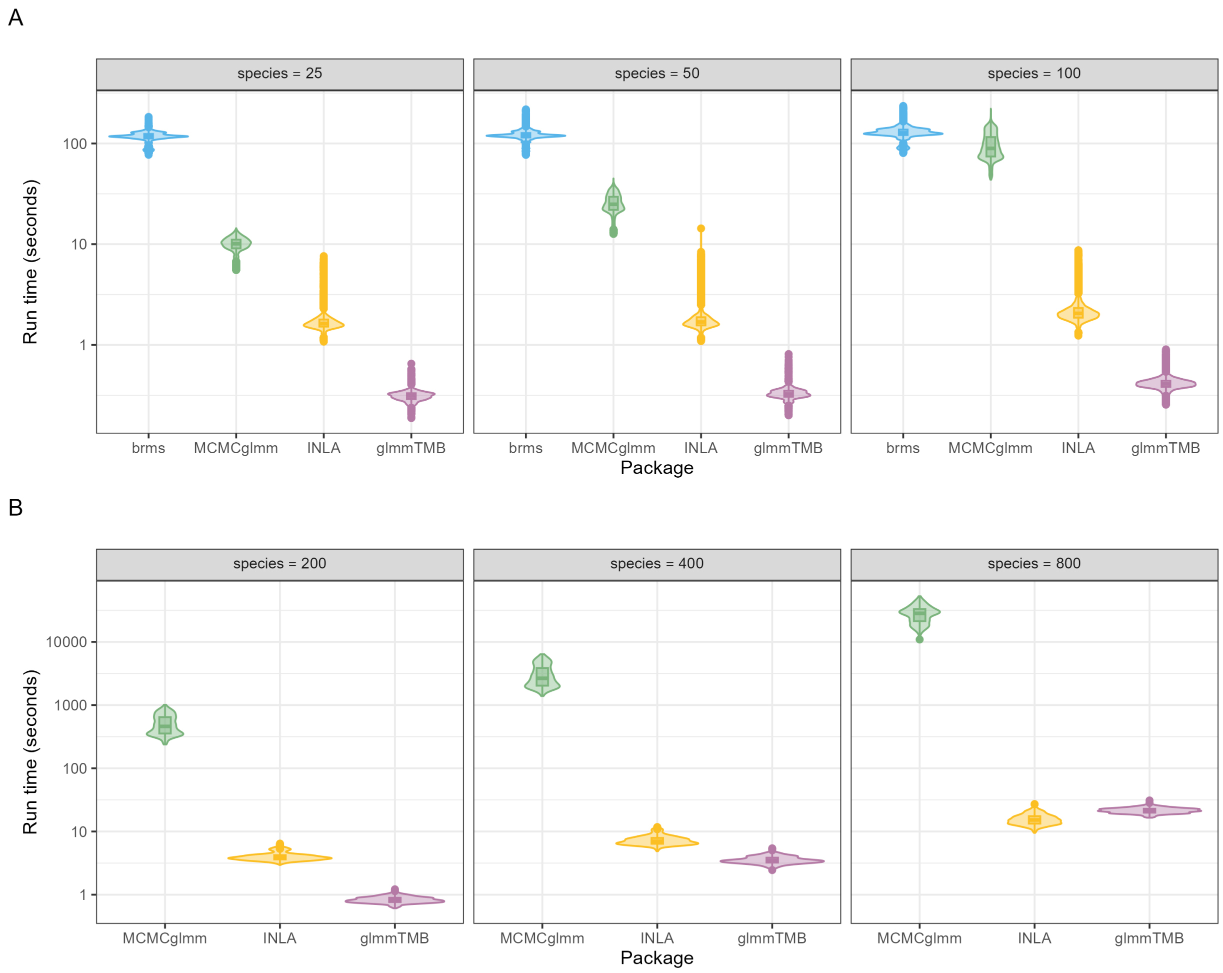

Figure S5: Run times (seconds) for phylogenetic generalised linear mixed models fitted using four R packages (`brms`, `MCMCglmm`, `INLA`, `glmmTMB`). The run time axis is displayed on a $\log_{10}$ scale for visualisation. Panels show (A) results from $\sim 30,000$ converged simulated models across datasets with 25 to 100 species, and (B) results from $\sim 3,000$ converged simulated models with 200 to 800 species. All models used the same fixed and random effects structure but different parameter values.

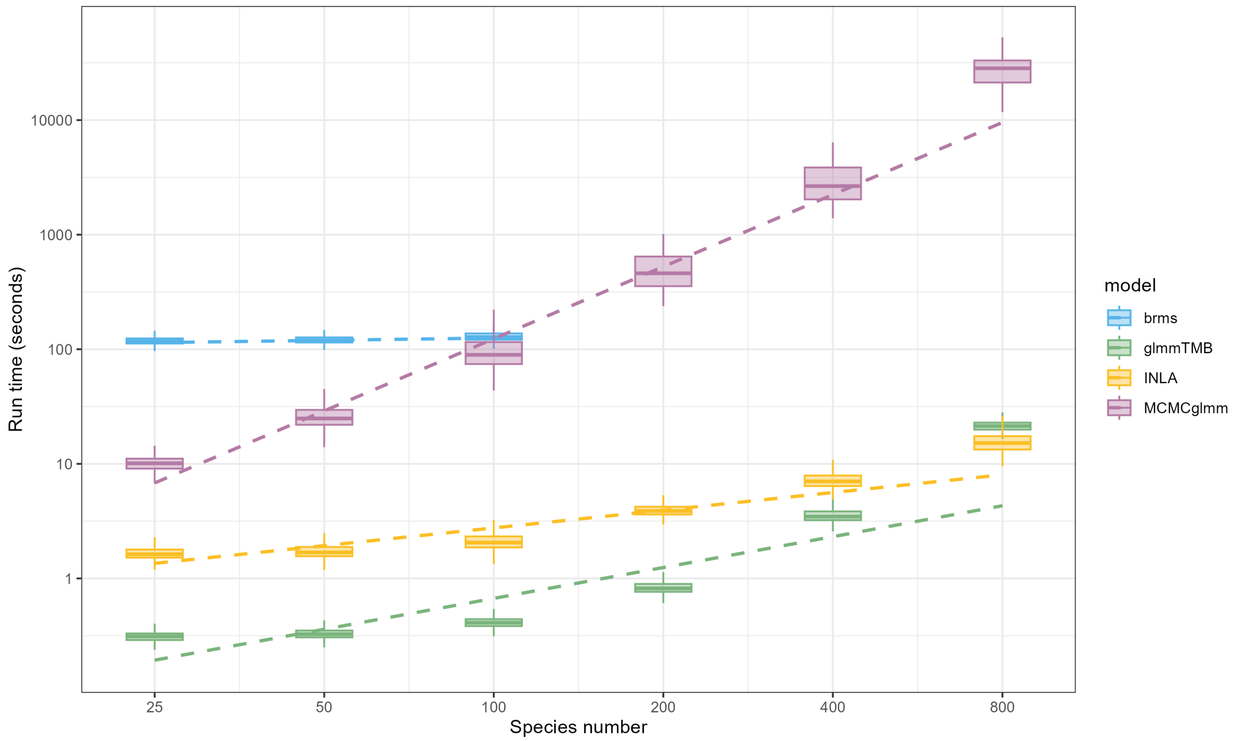

Figure S6: Computation times (seconds) for fitting phylogenetic models across packages in relation to species number, plotted on logarithmic axes. Boxplots show run time distributions per package, with coloured dashed lines indicating fitted regressions.

###### 2.2.3 Fixed covariate estimate

Table S10: Monte Carlo Standard Errors (MCSE) of the estimate $\hat{\beta}_1$ bias per package and per number of species.

| package | species size | $\hat{\beta}_1$ MCSE |
|---|---|---|
| INLA | 25 | 0.1042 |
| MCMCglmm | 25 | 0.1004 |
| brms | 25 | 0.1001 |
| glmmTMB | 25 | 0.1006 |
| INLA | 50 | 0.0716 |
| MCMCglmm | 50 | 0.0700 |
| brms | 50 | 0.0699 |
| glmmTMB | 50 | 0.0701 |
| INLA | 100 | 0.0483 |
| MCMCglmm | 100 | 0.0479 |
| brms | 100 | 0.0479 |
| glmmTMB | 100 | 0.0479 |
| INLA | 200 | 0.0333 |
| MCMCglmm | 200 | 0.0331 |
| glmmTMB | 200 | 0.0332 |
| INLA | 400 | 0.0228 |
| MCMCglmm | 400 | 0.0228 |
| glmmTMB | 400 | 0.0228 |
| INLA | 800 | 0.0158 |
| MCMCglmm | 800 | 0.0158 |
| glmmTMB | 800 | 0.0158 |

Table S11: Monte Carlo Standard Errors (MCSE) of $\hat{\beta}_1$ coverage rates per package and per number of species.

| package | species size | mean cov $\hat{\mu}$ | cov $\hat{\mu}$ MCSE |
|---|---:|---:|---:|
| brms | 25 | 0.961 | 0.1376 |
| MCMCglmm | 25 | 0.954 | 0.1478 |
| INLA | 25 | 0.935 | 0.1741 |
| glmmTMB | 25 | 0.936 | 0.1736 |
| brms | 50 | 0.949 | 0.1557 |
| MCMCglmm | 50 | 0.947 | 0.1589 |
| INLA | 50 | 0.939 | 0.1696 |
| glmmTMB | 50 | 0.938 | 0.1699 |
| brms | 100 | 0.950 | 0.1535 |
| MCMCglmm | 100 | 0.950 | 0.1539 |
| INLA | 100 | 0.947 | 0.1586 |
| glmmTMB | 100 | 0.947 | 0.1580 |
| MCMCglmm | 200 | 0.951 | 0.1533 |
| INLA | 200 | 0.945 | 0.1608 |
| glmmTMB | 200 | 0.949 | 0.1548 |
| MCMCglmm | 400 | 0.952 | 0.1517 |
| INLA | 400 | 0.951 | 0.1532 |
| glmmTMB | 400 | 0.950 | 0.1547 |
| MCMCglmm | 800 | 0.973 | 0.1147 |
| INLA | 800 | 0.971 | 0.1189 |
| glmmTMB | 800 | 0.973 | 0.1147 |

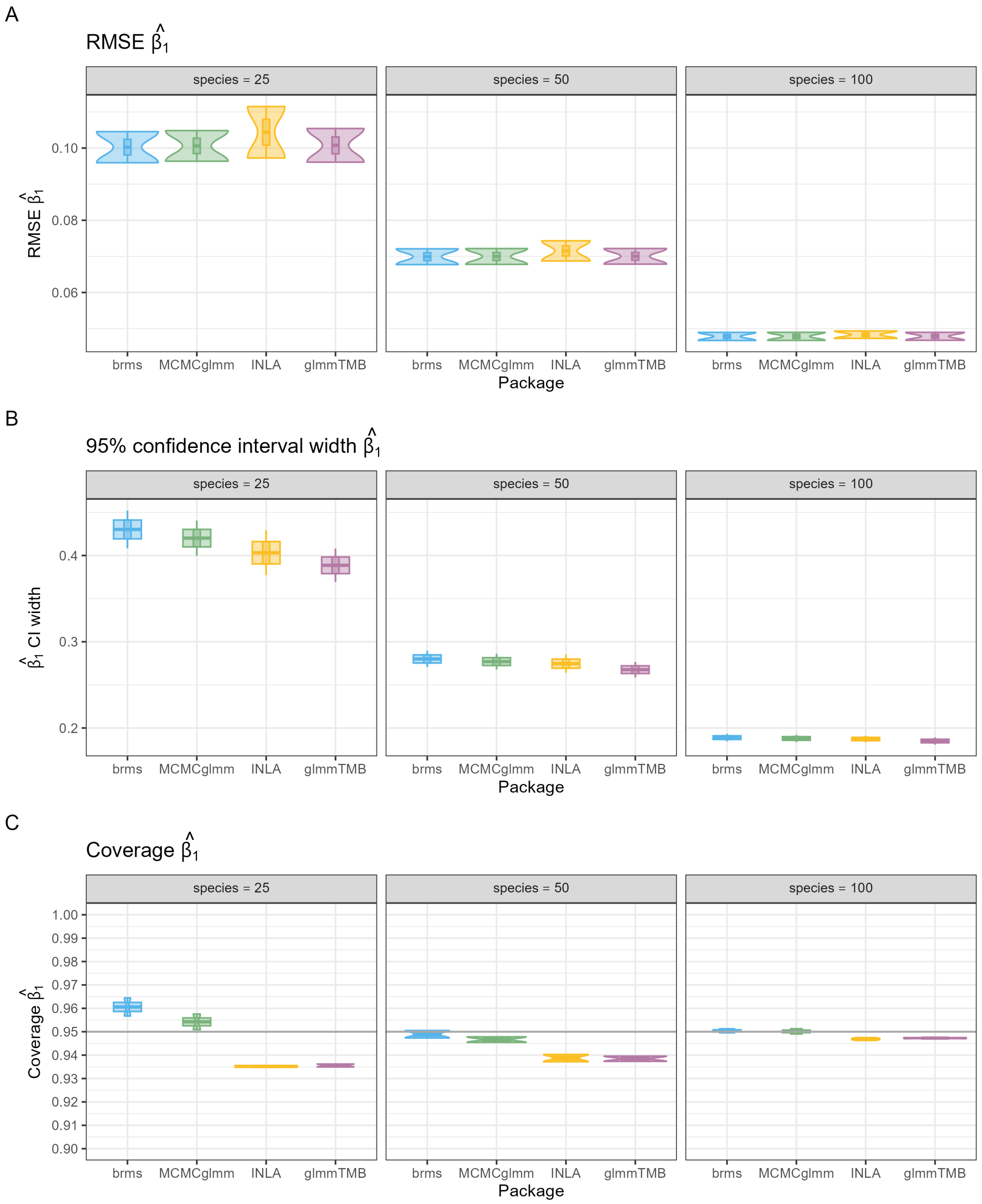

Figure S7: Comparison of five R packages in estimating the fixed effect $\hat{\beta}_1$ for datasets with 25, 50, and 100 species. Panels show (A) root-mean-square error (RMSE), (B) 95% confidence interval width, and (C) 95% interval coverage of the fixed effect estimate $\hat{\beta}_1$. Violin plots summarise simulation results across three conditions of $N_{\text{species}}$, and two conditions of $\hat{\sigma}_p^2$ ($3 \times 2 = 6$ conditions in total), each based on simulation replicates of converged models.

###### 2.2.4 Variance component estimates

Table S12: Monte Carlo Standard Errors (MCSE) of variance component ($\hat{\sigma}_p^2$) estimate bias per package and per number of species.

| package | species size | $\hat{\sigma}_p^2$ MCSE | $\hat{\sigma}_e^2$ MCSE |
|---|---|---|---|
| brms | 25 | 0.199 | 0.078 |
| MCMCglmm | 25 | 0.234 | 0.071 |
| INLA | 25 | 0.200 | 0.117 |
| glmmTMB | 25 | 0.172 | 0.074 |
| brms | 50 | 0.195 | 0.050 |
| MCMCglmm | 50 | 0.202 | 0.048 |
| INLA | 50 | 0.163 | 0.061 |
| glmmTMB | 50 | 0.170 | 0.048 |
| brms | 100 | 0.167 | 0.033 |
| MCMCglmm | 100 | 0.166 | 0.032 |
| INLA | 100 | 0.142 | 0.037 |
| glmmTMB | 100 | 0.150 | 0.032 |
| MCMCglmm | 200 | 0.144 | 0.021 |
| INLA | 200 | 0.130 | 0.022 |
| glmmTMB | 200 | 0.134 | 0.021 |
| MCMCglmm | 400 | 0.127 | 0.015 |
| INLA | 400 | 0.118 | 0.015 |
| glmmTMB | 400 | 0.122 | 0.015 |
| MCMCglmm | 800 | 0.120 | 0.011 |
| INLA | 800 | 0.113 | 0.011 |
| glmmTMB | 800 | 0.117 | 0.011 |

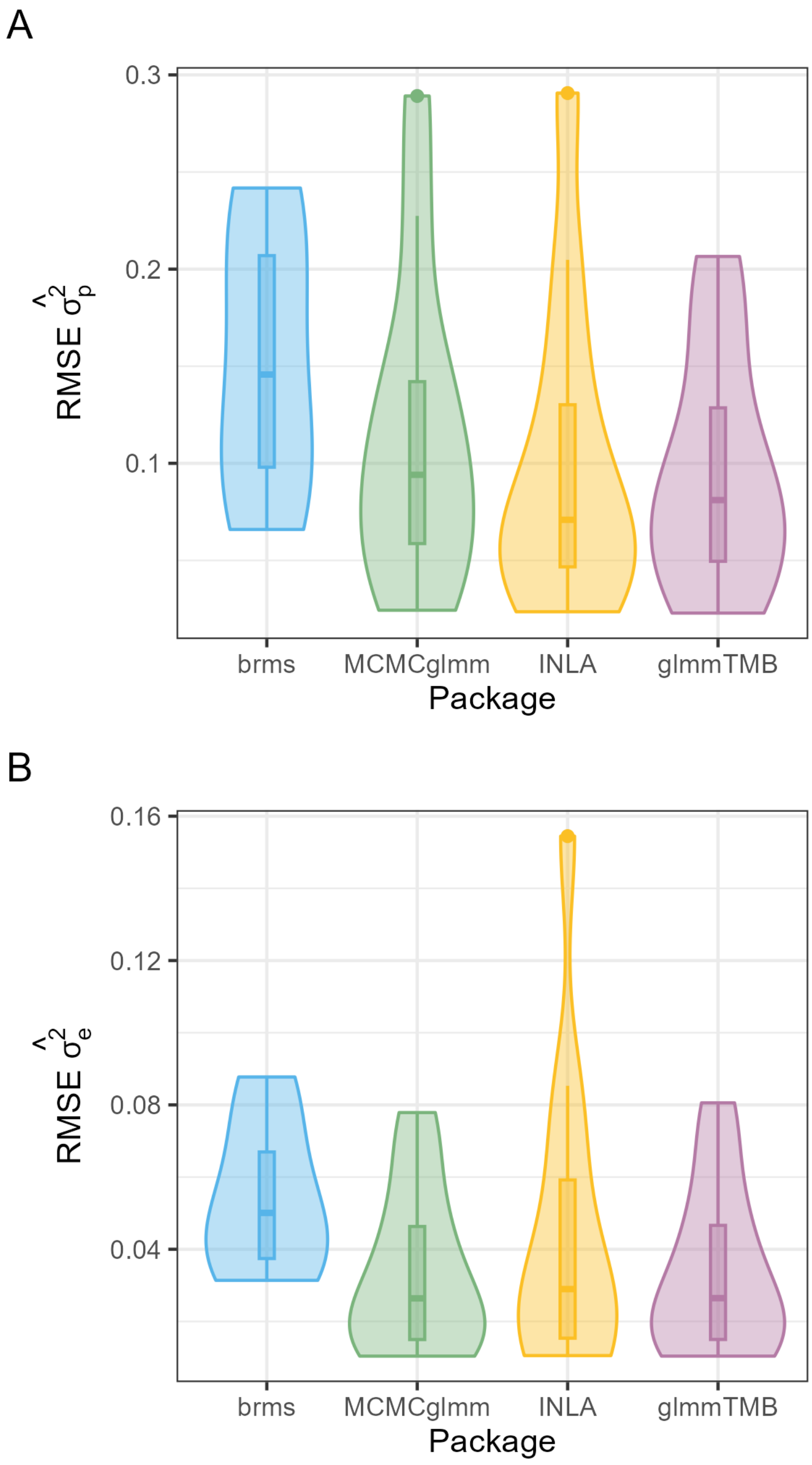

Figure S8: Root mean-squared error (RMSE) of variance component estimates across five R packages. Panels show measures of RMSE of (A) species-level variance $\hat{\sigma}^2_s$, (B) phylogenetic variance $\hat{\sigma}^2_p$, and (C) residual variance $\hat{\sigma}^2_e$. Violin plots summarise simulation results across all (6) conditions of $N_{\text{species}}$ and all (2) conditions of $\hat{\sigma}^2_p$, each based on simulation replicates of converged models.

#### 2.3 Case study 1: Bird colour evolution

##### 2.3.1 Continuous colour trait

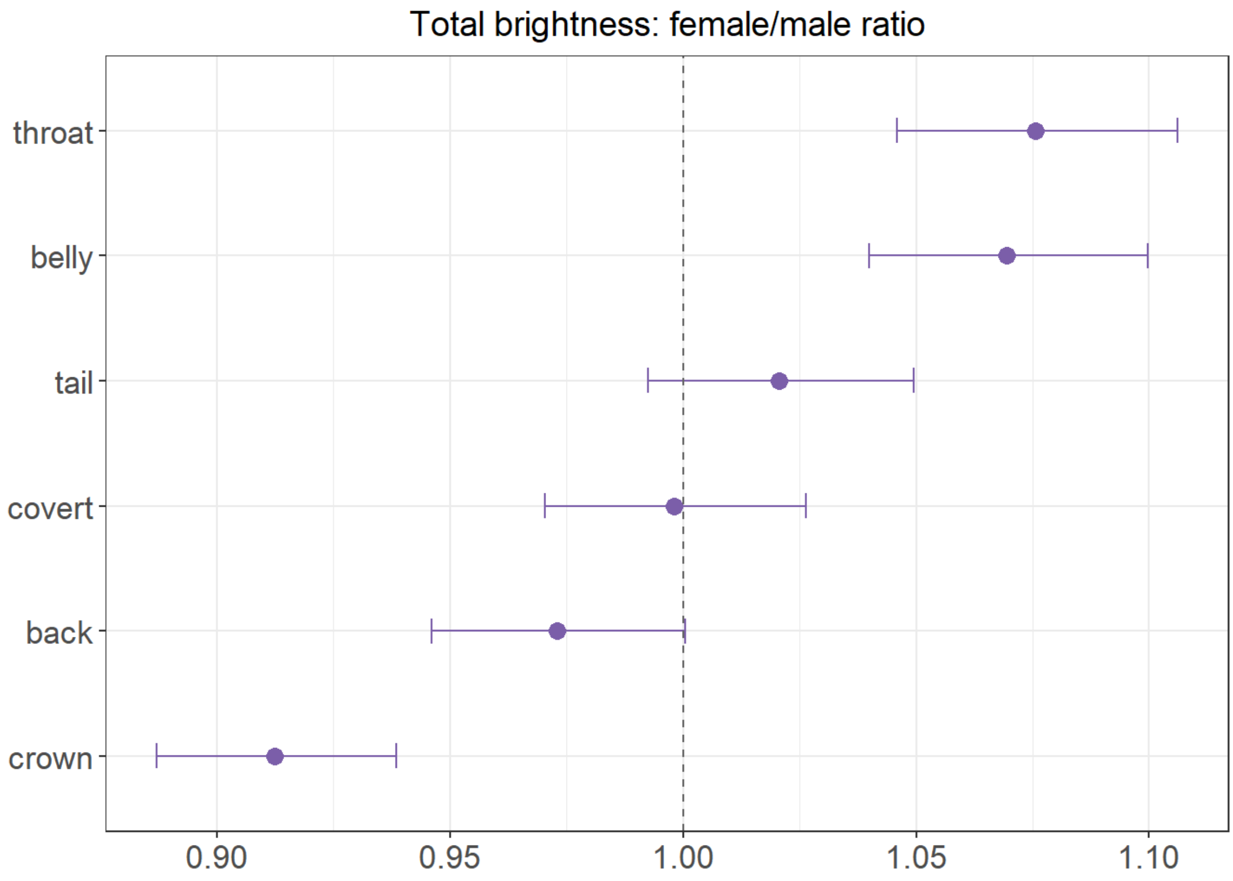

Figure S9: Female-to-male brightness ratios estimated from a Gamma GLMM with a log link for B1 across the six body regions. Points represent marginal means and error bars the 95% credible intervals. Ratios above one indicate higher female total brightness, ratios below one indicate higher male total brightness.

###### 2.3.2 Binary colour trait

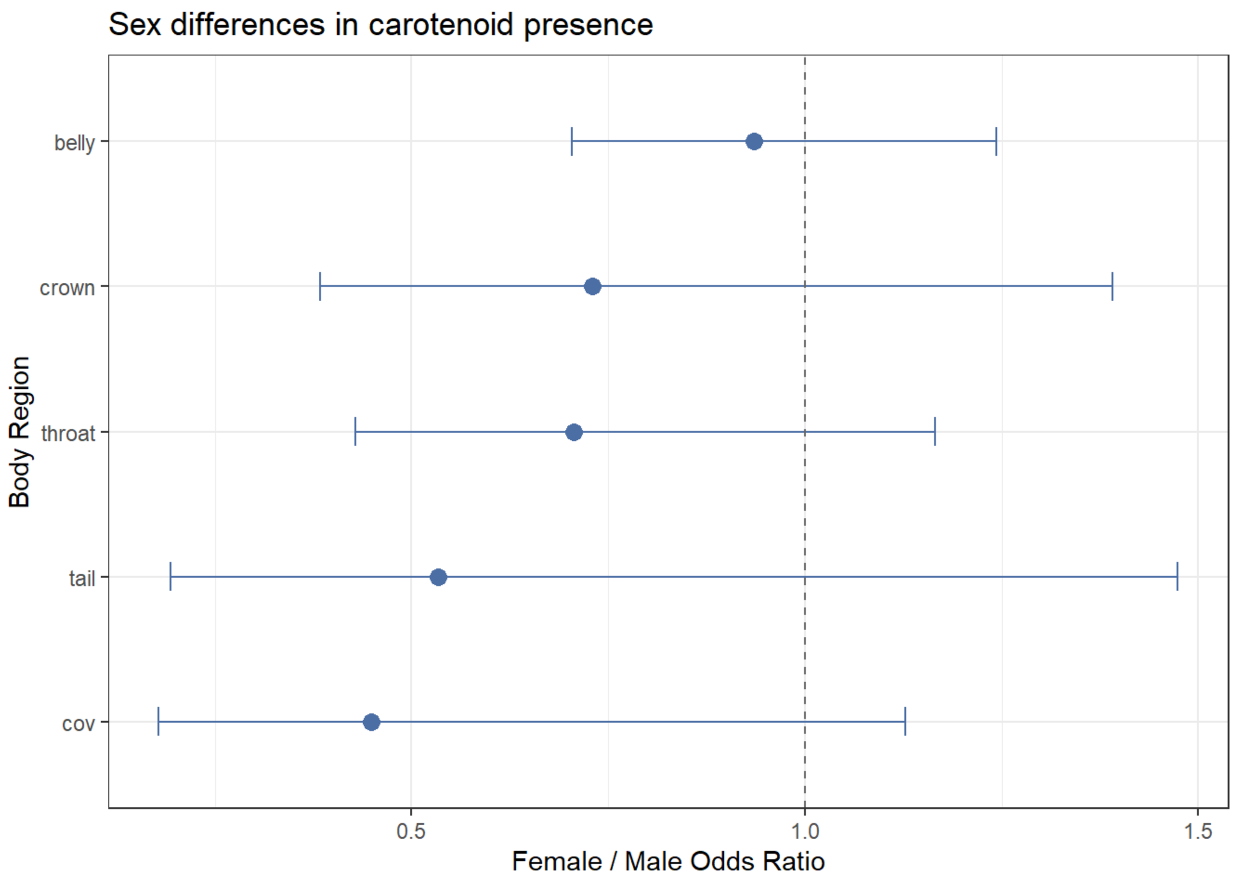

Figure S10: Female-to-male brightness ratios estimated from binomial GLMM with a logit link for presence of carotenoid color across the six body regions. Points represent marginal means and error bars the 95% credible intervals. Ratios above one indicate higher female total brightness, ratios below one indicate higher male total brightness.

###### 2.3.3 Ordinal colour trait

We analysed the ordered proportion $\texttt{prop\_carotenoid}_i$ of carotenoid colour across all body regions of individual birds using an ordinal beta GLMM, including a fixed effect of `sex`:

$$\texttt{prop\_carotenoid}_i \sim \text{Beta}(\mu_i, \phi)$$
$$\text{logit}(\mu_i) = \beta_0 + \beta_1 \; \texttt{sex}_i + s_{k[i]} + p_{k[i]}$$

where $\mu_i$ is the mean of the beta distribution on the logit scale, and $\phi$ is the precision parameter of the beta distribution.

```
glmmTMB(prop_carotenoid ~ sex + (1|species) + propto(0 + species|g,phylo.mat),
                   family = ordbeta(),
                   data = ord.dat)
```

We found that the model estimated that 96.7% of the total variance at the species-level random effects was explained by phylogeny, indicating a strong phylogenetic signal. The main contrast, expressed as the odds ratio of carotenoid presence between sexes, was significant (OR = 0.917, SE = 0.025, 95% CI: 0.870–0.966, z = –3.24, p = 0.0012), showing that males had a higher proportion of body regions with carotenoid presence than females.

##### 2.4 Case study 2: Macro-evolutionary patterns of hydraulic traits in plants

The second case study consisted of a published dataset of plant hydraulic traits Sanchez-Martinez et al., 2020 described in Figure S11. Hydraulic traits describe the ability of plants to transport water through their vascular system and to withstand water stress. These traits are fundamental to plant physiology, as they influence growth, survival, and adaptation to different environments.

###### 2.4.1 Hydraulic conductivity (kS)

Hydraulic conductivity ($k_s$) measures how efficiently water moves through the xylem of a plant stem. Species with high $k_s$ can transport water more rapidly, which supports fast growth but may also come with greater vulnerability to water stress. In contrast, species with lower $k_s$ transport water more slowly, reflecting a more conservative strategy.

We analysed the log hydraulic conductivity of each species using a Gaussian GLMM with REML, including a fixed effect of plant group and two genus-level random effects (unstructured and phylogenetic). The variable `group` is binary and indicates angiosperm or gymnosperm.

$$\log(\text{kS}_i) \sim \mathcal{N}(\mu_i, \; \sigma^2)$$

$$\mu_i = \beta_0 + \beta_1 \, \text{group}_i + s_{\text{genus}[i]} + p_{\text{genus}[i]}$$

where $\text{group}_i$ is a binary variable indicating angiosperm or gymnosperm, $s_{\text{genus}[i]}$ is the genus random intercept, and $p_{\text{genus}[i]}$ is the phylogenetic random effect based on the genus correlation matrix.

```
glmmTMB(log(Ks) ~ group + (1|genus) + propto(0 + genus|g, phylo.mat.p),
                      family = gaussian(),
                      REML=TRUE,
                      data = ks.dat)
```

For the hydraulic conductivity trait ($k_S$), the Gaussian PGLMM indicated that phylogenetic effects explained 83.9% of the total species-level variance, with the remaining 16.1% attributed to non-phylogenetic factors. Estimated marginal means on the response scale suggested that angiosperms had higher $k_S$ values (1.22, 95% CI [0.27, 5.51]) compared with gymnosperms (0.62, 95% CI [0.24, 1.64]). However, this difference was not statistically significant (ratio = 1.97, 95% CI [0.33, 11.8], $p = 0.46$).

### 2.4.2 P50

P50 represents the stem water potential at which a tree experiences a 50% loss of hydraulic conductivity. More negative values indicate greater resistance to hydraulic stress, meaning that species with lower P50 are more resilient to drought conditions (Trugman et al., 2020).

We analysed the negative of P50 (sign-flipped values) using a Gamma PGLMM, with a fixed effect of plant group (angiosperms vs gymnosperms), a genus random intercept, and a phylogenetic random effect.

$$-\mathrm{P50}_i \sim \mathrm{Gamma}(\mu_i,\ \phi)$$

$$\log(\mu_i) = \beta_0 + \beta_1\,\mathrm{group}_i + s_{\mathrm{genus}[i]} + p_{\mathrm{genus}[i]}$$

where $\mathrm{group}_i$ indicates angiosperm or gymnosperm, $s_{\mathrm{genus}[i]}$ is the genus random intercept, and $p_{\mathrm{genus}[i]}$ is the phylogenetic random effect based on the genus correlation matrix.

```
glmmTMB(negP50 ~ group + (1|genus) + propto(0 + genus|g,phylo.mat.p),
                      family = Gamma(link = "log"),
                      data = p50.dat)
```

For the trait($P_{50}$, the Gamma PGLMM indicated that phylogenetic effects explained 73.7% of the total species-level variance. Estimated marginal means on the response scale showed that gymnosperms had lower (more negative) $P_{50}$ values (4.24, 95% CI [2.38, 7.54]) compared with angiosperms (2.21, 95% CI [0.88, 5.53]). However, this difference was not statistically significant (ratio = 0.52, 95% CI [0.18, 1.54], $p = 0.24$).

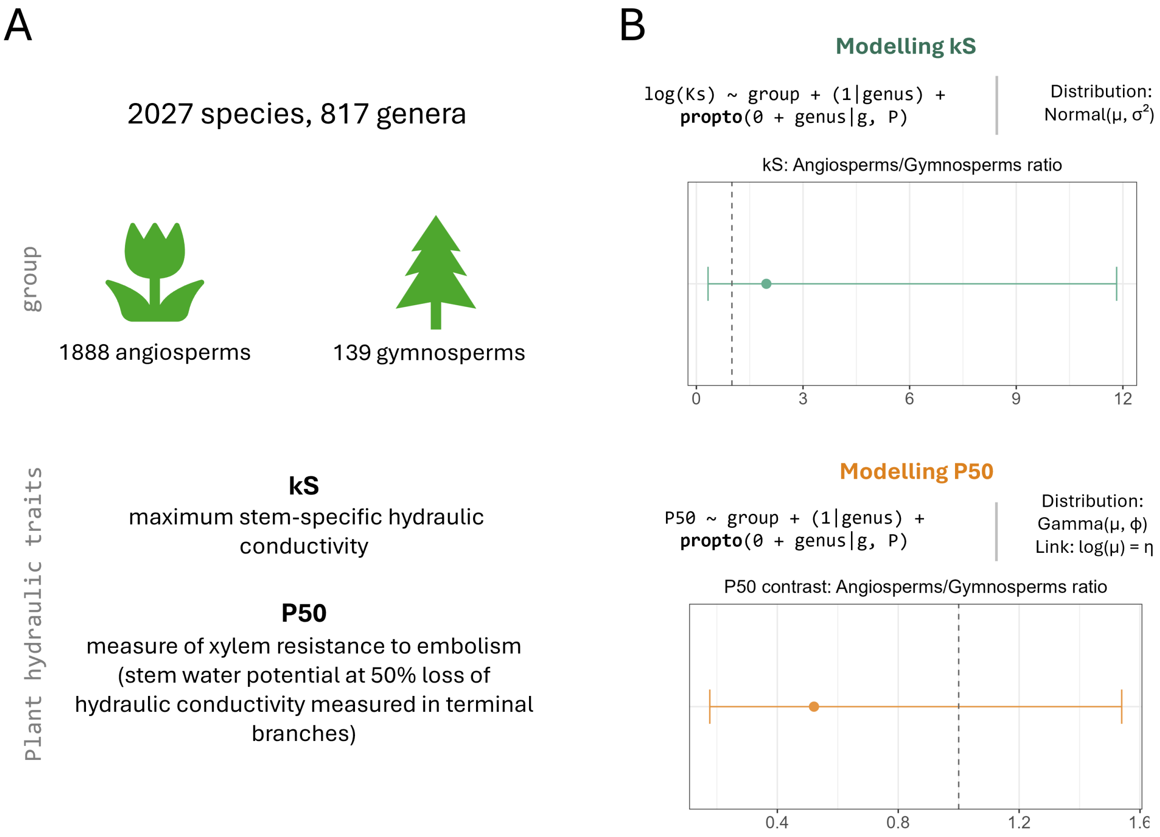

Figure S11: **Case study of global macro-evolutionary patterns of hydraulic traits in plants.**
**(A)** Overview of the published dataset Sanchez-Martinez et al., 2020, comprising 2027 species in total (1888 angiosperms and 139 gymnosperms). Two hydraulic traits were analysed: $k_s$, maximum stem-specific hydraulic conductivity, and P50, the stem water potential at 50% loss of hydraulic conductivity (measured in terminal branches) as an indicator of resistance to embolism.
**(B)** Estimated angiosperm-to-gymnosperm contrasts for each trait. Points indicate ratio estimates and error bars the 95% confidence intervals. Ratios above one indicate higher mean values in angiosperms, and ratios below one indicate higher mean values in gymnosperms.
